# Notch controls arterialization by regulating the cell cycle and not differentiation

**DOI:** 10.1101/2020.07.07.192344

**Authors:** Wen Luo, Irene Garcia-Gonzalez, Macarena Fernandez-Chacon, Veronica Casquero-Garcia, Rui Benedito

**Affiliations:** Molecular Genetics of Angiogenesis Group. Centro Nacional de Investigaciones Cardiovasculares (CNIC), Melchor Fernandez Almagro, 3, Madrid, Spain

## Abstract

Arteries are thought to be formed by the induction of a highly conserved arterial genetic program in a subset of vessels experiencing an increase in pulsatile and oxygenated blood flow. Both VEGF and Notch signalling have been shown to be essential for the initial steps of arterial specification. Here, we combined inducible genetic mosaics and transcriptomics to modulate and understand the function of these signalling pathways on cell proliferation, arterial-venous differentiation and mobilization. We observed that endothelial cells with high VEGF or Notch signalling are not genetically pre-determined and can form both arteries and veins. Importantly, cells completely lacking the Notch-Rbpj transcriptional activator complex can form arteries when the Myc-dependent metabolic and cell-cycle activity is suppressed. Thus, arterial development does not require the induction of a Notch-dependent arterial differentiation program, but rather the timely suppression of the endothelial metabolism and cell-cycle, a process preceding arterial mobilization and complete differentiation.

## Introduction

Angiogenesis occurs through the reiterative growth and remodeling of a precursor angiogenic vascular plexus. During the growth phase, endothelial cells (ECs) proliferate and migrate in order to expand the capillary network into hypoxic tissues rich in vascular endothelial growth factor (VEGF). After the formation of a rudimentary capillary plexus, the existing capillary branches require active remodeling to form arterial vessels specialized in oxygenated blood delivery and venous vessels specialized on low-oxygen blood collection and its return to the pumping heart. This stepwise process of angiogenesis and vascular remodeling involves numerous genes and signaling pathways ^1–5^, whose activity must be regulated with high spatiotemporal definition to ensure proper vascular development.

Experimental evidence obtained in diverse angiogenesis settings and model organisms has shown that the VEGF and Notch signalling pathways are essential regulators of angiogenesis. VEGF is a positive regulator of endothelial sprouting and proliferation, whereas Notch inhibits these processes ^6, 7^ in a manner dependent on the mitogenic context ^8^. These pathways also play key roles in the differentiation of progenitor and capillary ECs to fully defined arterial ECs ^1, 4, 7, 9–12^. Expression of the main arterial Notch ligand Delta-like 4 (Dll4) is induced by VEGF and is highest in arterial and sprouting tip ECs, being lower in capillaries and absent from most venous ECs ^7, 13, 14^. The primordial vessels of embryos lacking endothelial Vegfr2 or Dll4-Notch signalling fail to acquire features of arterial molecular identity, such as the expression of Efnb2, Cx37 and Cx40 ^9, 10, 15–19^. Conversely, increased expression of Dll4 results in the early expansion or dilation of anterior arterial vessels and the ectopic induction of arterial molecular markers in the venous endothelium ^20^.

Recent studies in mouse retinas and zebrafish have provided increased mechanistic insight into arterialization, showing that it involves the contraflow migration of pre-determined Cxcr4^High^/Notch^high^ ECs towards the developing arteries ^21, 22^. scRNAseq and lineage tracing in mouse hearts, similarly showed that pre-arterial ECs with higher Cxcr4, Cx40 and Notch signaling are committed to form the main coronary arteries ^23^, whereas full loss of Dll4/Notch signalling results in the loss of arterial EC identity and coronary arteries ^24, 25^. These results suggest that arterialization involves the genetic pre-determination of arterial identity within a remodelling capillary network ^1^, that is later refined and maintained by arterial blood flow ^4, 26, 27^.

Molecularly, activation of surface Notch receptors by its ligands triggers the release of the Notch intracellular domain (NICD) to the nucleus where it interacts with the cofactor Rbpj to promote a multitude of cell-type specific and context-dependent gene expression and cell functions, usually related to cell proliferation or differentiation ^28^. The current view is that the Notch-Rbpj transcriptional activator complex is a master regulator of arterial cell differentiation ^1, 4, 9, 15, 17, 29, 30^. However, Notch also actively inhibits the cell cycle of ECs, even before they form arteries ^7, 8, 12, 31^. Interestingly, arterial development coincides with the suppression of the cell cycle in ECs experiencing higher pulsatile and oxygenated blood flow ^23, 27, 32^. The sum of this evidence led us to question if the primary effect of Notch-Rbpj on arterialization is on cell differentiation, or on metabolic and cell-cycle regulation.

Here, we use multispectral genetic mosaics and combinatorial conditional genetics to define at high spatiotemporal resolution the role of VEGF, Notch and Myc in the regulation of the cell-cycle and arteriovenous differentiation of coronary ECs. This approach revealed that both high VEGF signalling and high Notch signalling prime ECs to form arteries by suppressing Myc-dependent mitogenic activity in coronary pre-arterial ECs. When located in a pre-venous microenvironment, ECs with high VEGF and Notch signalling can also form veins, suggesting that they are not genetically pre-determined to arterial EC fate. Remarkably, ECs can acquire arterial markers and identity in the full absence of Notch–Rbpj transcriptional activity if the Myc-dependent cell cycle and metabolic activity are suppressed. Our study suggests that the primary effect of the Notch–Rbpj transcriptional machinery is to inhibit the cell cycle and metabolism of ECs located in pre-arterial capillaries and that full arterial cell differentiation is a subsequent step likely involving the integration of additional genetic and biophysical factors.

## Results

### Genetic mosaics reveal the fate of heart ECs with distinct Notch signalling levels

To determine how normal coronary ECs clonally expand and migrate over time, we crossed *Dual ifgMosaic* mice ^33^ with *Tie2-Cre* mice ^34^ and analysed the clonal distribution of wildtype ECs with distinct multispectral barcodes in the heart vasculature. As shown before during embryoid body and mouse retina angiogenesis ^33, 35^, coronary ECs intermix extensively during their migration and proliferation giving rise to highly mosaic heart vessels (Fig. 1a and Extended Data Fig. 1a,b).

**Fig. 1.**
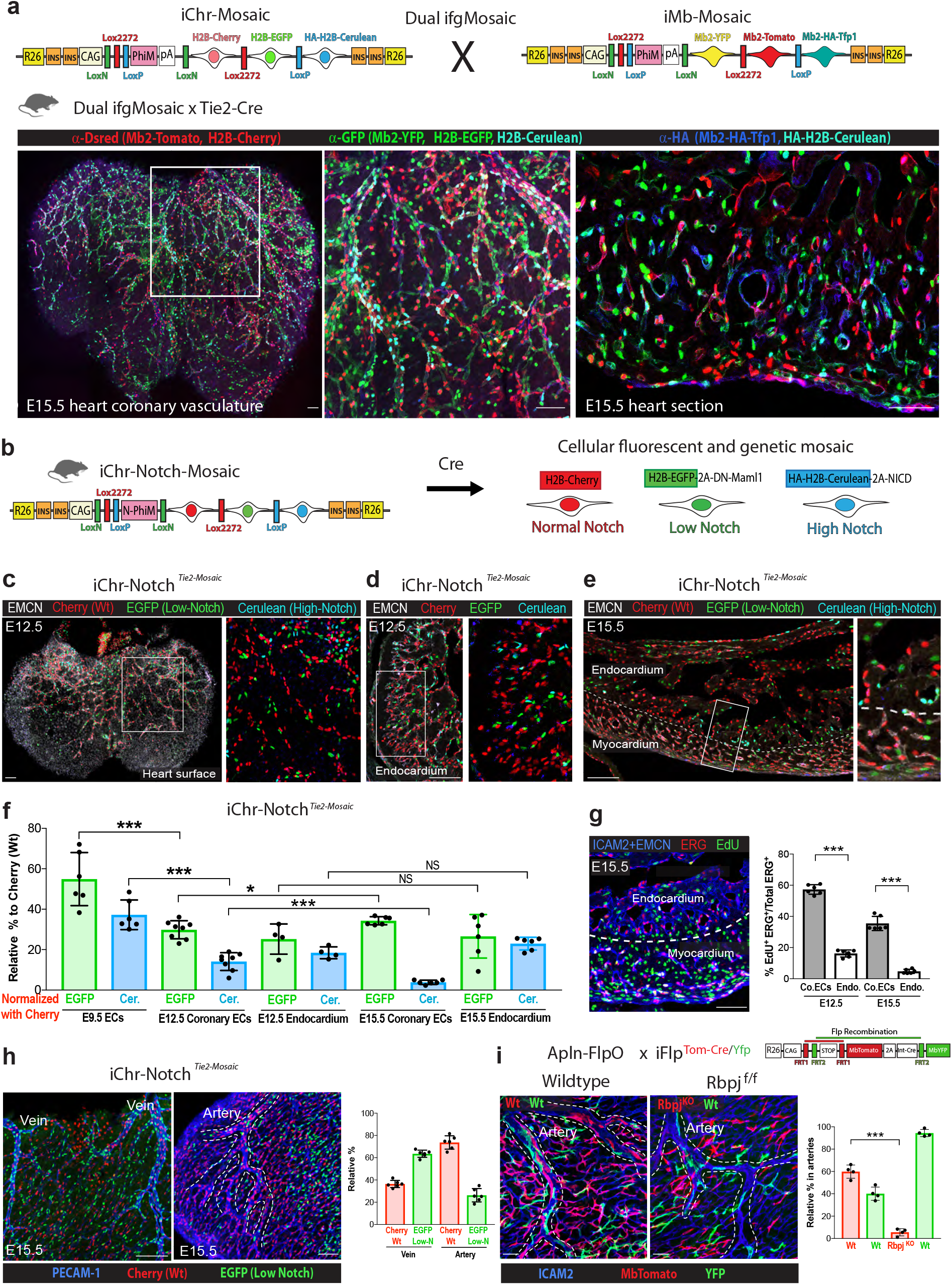
Fate mapping of heart ECs having distinct Notch signalling levels. (a) Schematic representation of Dual ifgMosaic constructs used to fate map wildtype coronary ECs with high multispectral clonal resolution (see also Extended Data Fig. 1a,b). (b) Mice having the iChr-Notch-Mosaic and Tie2-Cre alleles undergo induction of a multispectral mosaic of ECs expressing distinct Notch signalling modulators and having distinct Notch levels (see also Extended Data Fig.1c). (c-e) Confocal images of whole embryonic hearts or sections showing the localization and relative frequency of ECs having distinct Notch signalling levels. (f) Quantification of c, d, e showing how the relative percentage of each cell type changes throughout development (minimum n=3 hearts per stage). (g) Representative confocal image showing analysis of EC proliferation (ICAM2+EMCN+/ERG+/EdU+) in the myocardium and endocardium. Quantification at E12.5 and E15.5 (n=3 hearts per stage). (h) Contribution of ECs with distinct Notch signalling levels to the development of coronary veins or arteries (n=3 hearts per group). (i) Mice carrying the alleles Apln-FlpO and iFlp^MbTomato-Cre/MYfp^ have induction of an EC mosaic in growing coronary vessels. When these alleles are on a wildtype background, both MbTomato and MbYFP label wildtype cells. When these alleles are on a Rbpj^fl/fl^ background only Tomato-2A-Cre+ cells have Rbpj deletion (see also Extended Data Fig. 1j,k). Quantification shows that ECs with Rbpj deletion rarely form arteries (n=4 hearts per group). Scale bars, 100 um. Error bars indicate SD. *p < 0.05, ***p < 0.001. NS, non significant.

To map the clonal expansion and arterio-venous fate of single ECs with permanently decreased or increased Notch signalling, we induced loss and gain-of-function genetic mosaics by crossing *iChr-Notch-Mosaic* mice ^8, 33^ with *Tie2-Cre* mice. This induces a mosaic of cells at embryonic day (E) 8.5 having normal (H2B-Cherry+), low (H2B-GFP+) or high (HA-H2B-Cerulean+) Notch signalling (Fig. 1b and Extended Data Fig.1c) throughout the embryonic endothelium (Tie2-Cre+). In contrast to global or EC-specific *Notch1, Dll4 or Rbpj* mutants, *iChr-Notch*^*Tie2-Mosaic*^ mosaic embryos developed normally and survived to adulthood (Extended Data Fig.1d). This allowed us to compare the proliferative, migratory and differentiation dynamics of the different mutant cells with neighboring control/wildtype (Cherry+) cells sharing the same environment. ECs with high Notch signalling (Cerulean+) were strongly outcompeted during coronary vessel development (Fig. 1c-f, compare E9.5 with E12.5 and E15.5 coronary ECs ratios). These results are consistent with the known lower proliferative activity of ECs with higher Notch signalling ^8, 36–39^. However, unlike the situation in the coronary vessels, higher Notch activity did not generate a proliferative disadvantage in the endocardium (Fig. 1c-f, compare the relative % of Cerulean+ cells in E12.5 and E15.5 endocardium). Endocardial cells are significantly less proliferative than coronary ECs (Fig. 1g), which may explain why cells with high Notch signaling remain prevalent in the E15.5 endocardium upon pulsing with Tie2-Cre (pan-endothelial) or the Nfatc1-Cre (endocardium-specific) alleles (Fig. 1f and Extended Data Fig. 1e-g). Interestingly, coronary ECs with lower Notch signalling (GFP+) on average did not proliferate more than the adjacent wildtype (Cherry+) ECs (Fig. 1c-f and Extended Data Fig. 1h,i). We recently showed that ECs have a bell-shaped response to mitogenic stimuli and that cells with low Notch signalling can either proliferate more, or become arrested when the levels of VEGF and ERK activity are high ^8^. The developing myocardium is known to be rich in VEGF, which may explain why the GFP+ cells generally do not overproliferate and outcompete Cherry+ cells in this tissue.

Besides the regulation of EC proliferation, Notch has also been shown to induce arterial differentiation and mobilization of ECs to arteries ^1, 4, 7, 9–12^. In agreement with previous studies in zebrafish, we found that in *iChr-Notch*^*Tie2-Mosaic*^ hearts, ECs with a decrease in Notch signalling (GFP+), were more frequently found in veins, whereas ECs with normal Notch signalling (Cherry+) were more frequently found in arteries, however Notch low (GFP+) cells could be found in both types of vessels (Fig. 1h). These results indicate that the individual cell level of Notch signalling, influences its contribution to the formation of arterial or venous vessels but is not absolutely deterministic. Since the decrease in Notch signaling in ECs expressing DN-Maml1 is less pronounced on average than after full deletion of *Rbpj* ^8^, we also developed an alternative Flp-inducible genetic mosaic strategy (*iFlp*^*Tom-Cre/Yfp*^ *Mosaic*) to analyse the fate of single coronary ECs with (MbTomato-2A-Cre+) or without (MbYFP+) full deletion of *Rbpj* (Extended Data Fig. 1j). The transcription factor Rbpj is essential for inducing the Notch transcriptional program ^40^. In coronary vessel ECs expressing the X-chromosome-located *Apln-FlpO* allele (mosaic expression in female embryos due to random X-inactivation), a mosaic of Tomato-2A-Cre+ and YFP+ cells is generated at the coronary angiogenic front that will later form arteries. As expected, when in a *Rbpj floxed (Rbpj*^*f/f*^) background MbTomato-2A-Cre+ ECs showed deletion of *Rbpj* (*Rbpj*^*KO*^), whereas YFP+ cells expressed the gene normally (Extended Data Fig. 1k). Quantification of the relative distribution of these cells in control (*Rbpj*^*wt/wt*^) and mutant (*Rbpj*^*f/f*^) hearts revealed that cells full *Rbpj* deletion have a much more pronounced defect in artery formation than cells expressing DN-Maml1 (compare Fig. 1h with Fig. 1i). The relatively high frequency of individual *Rbpj*-negative ECs in periarterial capillaries indicates that these cells can locate in the proper pre-arterial capillary region but may fail to acquire the necessary pre-arterial phenotype to form or mobilize into developing coronary arteries. Alternatively, they might be excluded from developing arterial vessels with high Notch signaling.

### Ectopic Notch activation is compatible with vein development and impairs coronary artery development

To investigate the phenotypic consequence of increased endothelial Notch activity in all coronary vessels, we generated a new mouse line (*ROSA26*^*LSL-MbTomato-2A-H2B-GFP-2A-N1ICDP*^) that allowed us to induce fluorescent labelling of the membrane (MbTomato) and nuclei (H2B-GFP) of cells expressing the active Notch1 intracellular domain containing its native PEST sequence (*N1ICDP*), which results in a more moderate and physiologically relevant increase in Notch signalling ^41^ than achieved with other mouse lines ^42, 43^ (Fig. 2a,b). Consistent with this, embryos containing this allele and a *Tie2-Cre* allele (here abbreviated as, *N1ICDP*^*Tie2-Cre*^) die only after E15.5 (Extended Data Fig. 2a,b), rather than E9.5. This allowed us to study the phenotype of vessels in which a high proportion of ECs experience a physiologically relevant increase in Notch signaling during cardiovascular development from E8.5 to E15.5.

**Fig. 2.**
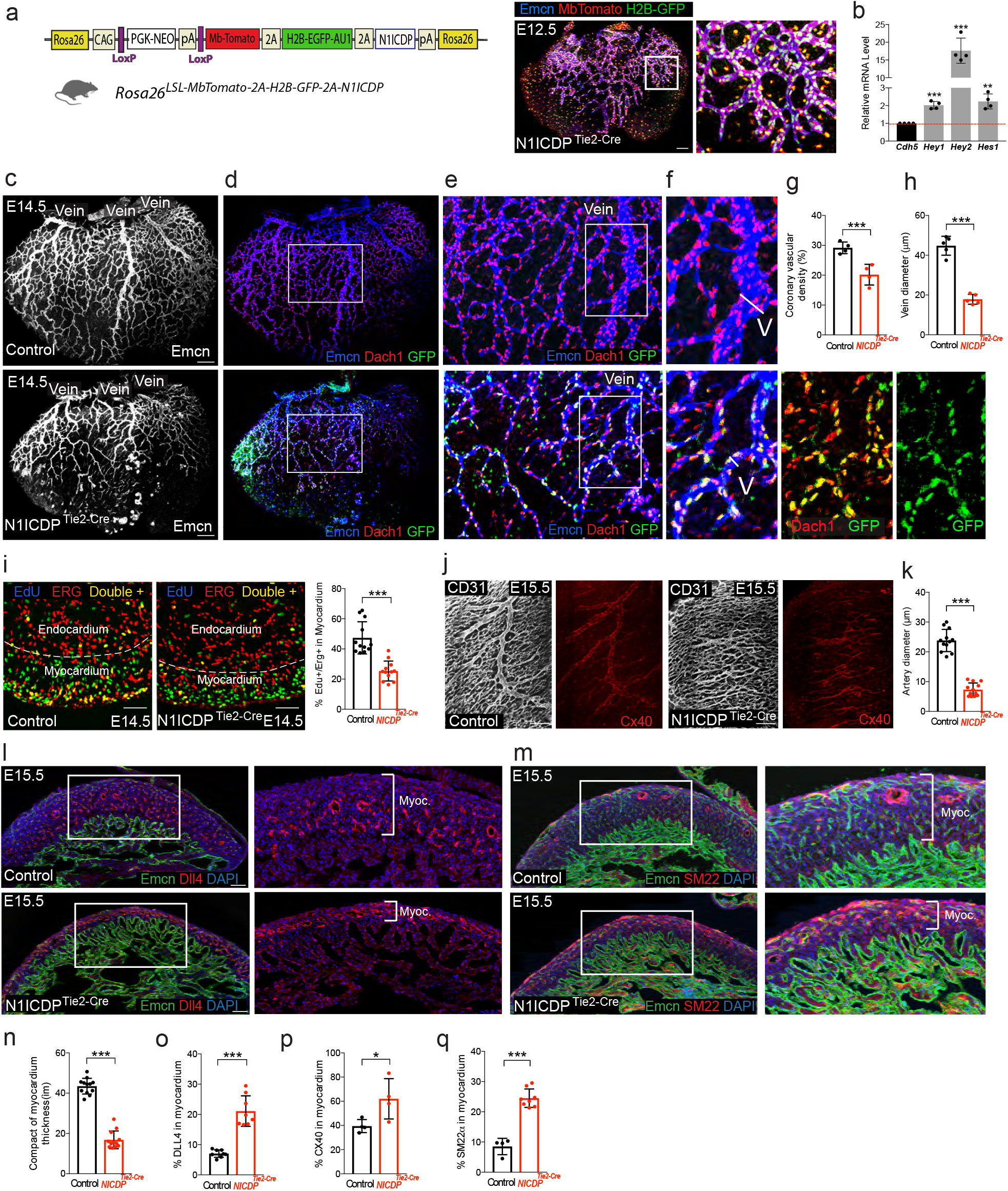
Ectopic Notch activation is compatible with vein development and impairs coronary artery development. (a) Genetic construct used to generate mice with Cre-dependent conditional expression of MbTomato, H2B-GFP and N1ICDP in Tie2-Cre+/Emcn+ coronary vessels. (b) Quantitative real-time PCR analysis of Notch target genes expression in CD31+/MbTomato+ and CD31+/MbTomato-cells isolated from mutant hearts (n=4 pooled litters). (c-h) Wholemount analysis of the heart surface coronary venous plexus in control and N1ICDP^Tie2-Cre^ mice showing a decrease in capillary density and larger vein (V) diameter (minimum n=4 per group). (i) Myocardium EC proliferation (ERG+/EdU+) analysis in sections of control and mutant hearts (n=3 per group). (j and k) Wholemount analysis of coronary arteries (CD31+/CX40+) in control and N1ICDP^Tie2-Cre^ mice (n=3 hearts per group). (l and m) Immunostaining of heart sections showing the endocardium (Emcn+) and the expression of arterial markers (Dll4 or SM22) in the compact myocardium (brackets inset) of control and mutant hearts. (n-q) Quantification of experiments represented in l and m and in Extended Data Fig. 2 (minimum n=4 hearts per group). Scale bars, 100 um. Error bars indicate SD. *p < 0.05, **p < 0.01, ***p < 0.001.

The moderate increase in EC Notch activity significantly inhibited the growth of coronary vessels (Fig. 2c-h), compromising myocardial development (Extended Data Fig. 2c,d). Despite their high Notch activity, GFP+ (N1ICDP+) ECs were frequently found in larger coronary veins; had lost the arterial marker Cx40 (Extended Data Fig. 2e) and had high expression of Endomucin and Dach1 (Fig. 2e,f), genes expressed by venous ECs. The mutant veins in *N1ICDP*^*Tie2-Cre*^ hearts were thinner and underdeveloped (Fig. 2f,h), presumably due to the lower proliferation rate of the mutant ECs in these embryos (Fig. 2i). Contrary to the prevailing view on the role of Notch signalling in arteriovenous specification, these results show that although coronary ECs with high Notch activity are biased to form arteries, these cells are not fully committed to the arterial fate since they are able to acquire venous identity markers and form veins. *N1ICDP*^*Tie2-Cre*^ veins have less ECs and are significantly thinner; however, this is an indirect consequence of the reduced proliferation of capillary ECs with high Notch activity, not to defects in capillary-to-venous differentiation or migration.

Increases in Notch activity have been previousl linked to an increase in arterialization ^9, 20^ and scRNAseq data revealed that pre-arterial ECs with higher *Cx40* expression and Notch signaling form the main coronary heart arteries ^23^. Surprisingly, *N1ICDP*^*Tie2-Cre*^ main coronary arteries were thinner and underdeveloped. The mutant hearts instead showed increased formation of small-diameter secondary collateral arteries in the region where the main coronary artery should form (Fig. 2j,k). These results show that ectopic Notch activation and Cx40 expression in pre-arterial capillaries impairs the development of larger coronary arteries, presumably by promoting the arterial phenotype in ectopic pre-arterial capillary branches that later divert blood flow from a single main artery, compromising its further development. Ectopic induction of the arterial phenotype in *N1ICDP*^*Tie2-Cre*^ coronary periarterial capillaries was evident from widespread and elevated expression of the arterial endothelial markers Cx40 and Dll4 (Fig. 2j,l,o,p and Extended Data Fig. 2f) and the increase in SM22alpha+ cells covering *N1ICDP*^*Tie2-Cre*^ capillaries (Fig. 2m,q).

We also crossed *Rosa26*^*ifN1ICDP*^ mice with *Nfact1-Cre* and *Pdgfb-CreERT2* mice, to determine the effect of Notch activation specifically in the endocardium (Nfatc1+) or in myocardial coronary vessels (Pdgfb+). Increase of Notch signaling in the endocardium was nonlethal and induced no major defects in cardiovascular development (Extended Data Fig. 2g-k). Adult *N1ICDP*^*Nfatc1-Cre*^ hearts were enlarged but showed no significant alteration in heart function or physiology (Extended Data Fig. 2l-n). In contrast, induction of Notch activity in coronary vessels with the *Pdgfb-CreERT2* allele at E11.5 and E12.5 produced similar cardiovascular defects to those of *N1ICDP*^*Tie2-Cre*^ embryos at E15.5 (Extended Data Fig.3). These results indicate that the cardiovascular development defects in *N1ICDP*^*Tie2-Cre*^ hearts result mainly from the activation of Notch signalling specifically in coronary vessels between E12.5 and E15.5.

### Coronary Dll4/Notch signalling suppresses a pro-mitogenic genetic program, not endothelial sprouting

To investigate the molecular mechanisms regulated by Dll4/Notch signalling specifically in coronary ECs, we isolated Pdgfb-IRES-GFP+ coronary ECs from *Control*^*Pdgfb-24h*^ and *Dll4*^*f/f-iDEC-Pdgfb-24h*^ E14.5 hearts. The transgene PAC *Tg(Pdgfb)-CreERT2*-*IresEGFP* ^44^ is only expressed in heart coronary ECs of the myocardium, and not in endocardial cells (Fig. 3a). In these embryos, CreERT2 activity was induced with tamoxifen at E13.5, 24h before collecting the cells for RNA extraction, at E14.5. This is the period pre-arterial coronary ECs become specified to form the first coronary arteries ^23^. *Dll4* genetic deletion for 24h resulted in complete downregulation of DLL4 protein and the Notch activity markers, active NICD, Cx40, *Hey1* and *Hes1* (Fig. 3b-d). RNAseq analysis revealed that full loss of Dll4-Notch signalling significantly alters the expression of 3360 genes (Fig. 3e). The set of differentially expressed genes (DEG) included downregulation of most coronary arterial-specific genes ^23^, whereas few coronary-vein-specific genes were upregulated (Fig. 3f), indicating that Dll4-Notch signalling is mainly essential for induction of the coronary arterial program, as also suggested by findings in other model organisms and systems ^9, 15–17^. Besides the significant loss of arterial identity marker genes, ECs with loss of Dll4-Notch signalling had a significant upregulation of genes related with cell proliferation and metabolism (E2F targets, G2M Checkpoint, Myc targets, Mtorc1 and glycolysis), which are linked to hyperactivated or hyperproliferative endothelial status (Fig. 3g,h and Extended Data Fig. 4a). ECs with loss of Dll4 also showed a significant increase in hypoxia marker genes (Extended Data Fig. 4a, b). Overall, these data show that the normal Dll4-Notch-induced arterialization is accompanied by a decrease in molecular signatures related to proliferation and metabolic activity.

**Fig. 3.**
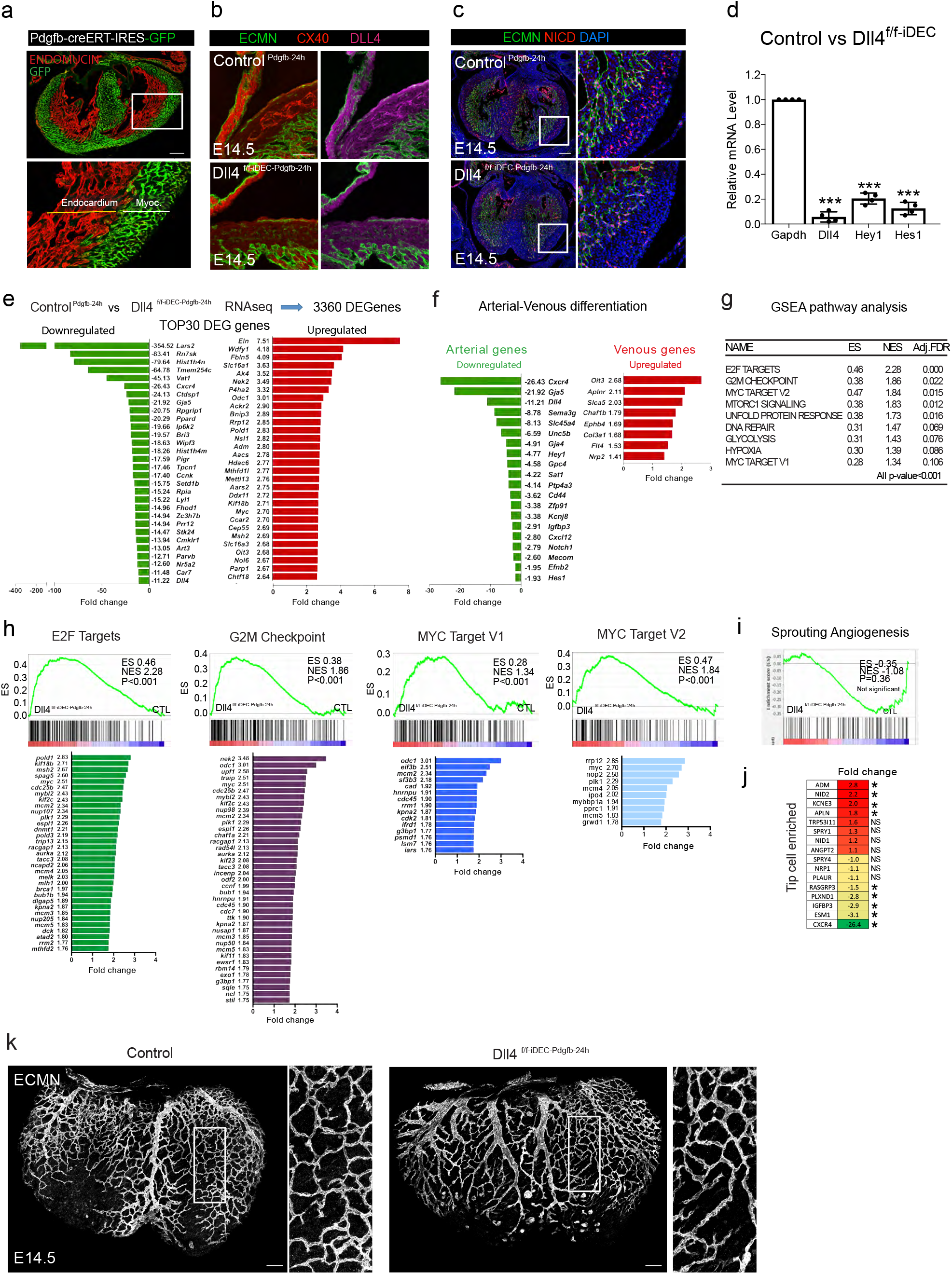
Transcriptional and phenotypic changes after loss of Dll4/Notch signalling in coronary ECs. (a) Confocal images of heart sections showing expression of the Pdgfb-CreERT2-IresEGFP allele only in myocardial vessels. (b-c) Immunostaining for Dll4, Cx40 and active N1ICD confirms the genetic deletion of Dll4 specifically in myocardial vessels and the significant downregulation of Notch signalling (active N1ICD) and arterial (Cx40) development. (d) qRT-PCR analysis of Pdgfb-CreERT2-IresEGFP+ myocardial ECs collected from wildtype and Dll4f/f hearts confirms the significant deletion of Dll4 and the downregulation of the Notch signalling target genes Hey1 and Hes1 (n=3 pooled litters per group, 2 replicates). (e) List of the 30 most up- and downregulated genes (from a total of 3360) after Dll4 deletion in coronary ECs for 24h. (f) List of the most significantly deregulated arterial and venous marker genes. (g) List of the top deregulated gene sets and their scores (GSEA MSigDB). (h) Complete list of deregulated genes within a selected enriched gene set (GSEA MSigDB). (i) Sprouting angiogenesis gene set is not significantly deregulated. (j) Fold change of tip-cell enriched genes (retina) in heart myocardial vessels after Dll4 deletion for 24h. (k) Confocal images of control and mutant heart surface vessels after loss of Dll4 expression for 24h. Scale bars, 100 um. Error bars indicate SD. *p < 0.05, **p < 0.01, NS, non-significant.

**Fig. 4.**
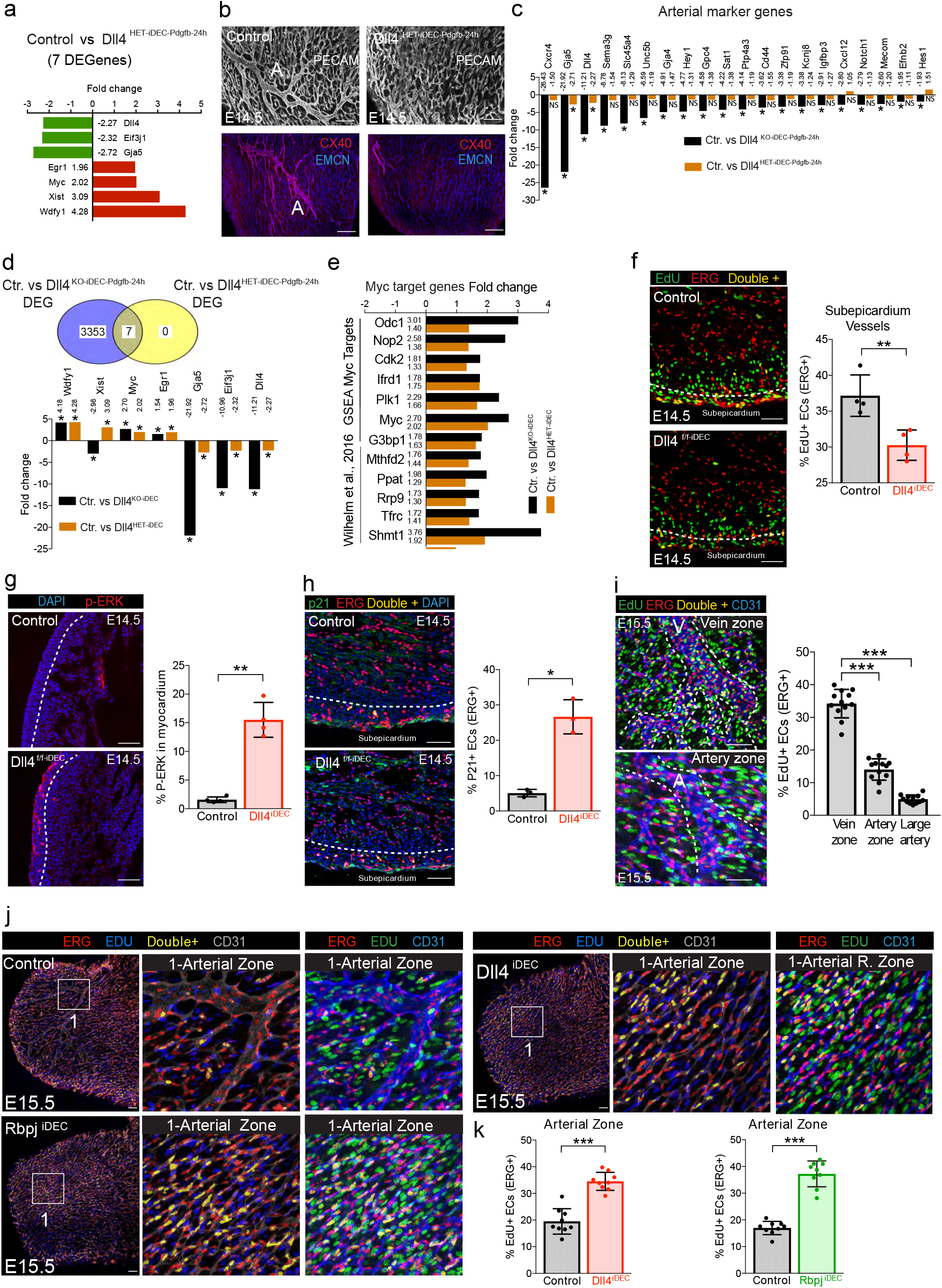
Metabolic and cell proliferation pathways are activated after hemizygous or complete loss of Dll4/Notch signalling. (a) List and fold change for the only 7 genes found to be significantly deregulated in Dll4 hemizygous mutant ECs (adjusted p-value < 0.05). (b) Hemizygous loss of Dll4 expression and function in myocardial ECs from E13.5 to E14.5 leads to lack of arterialization (CX40+). (c) List of arterial marker genes significantly downregulated (*) in myocardial ECs with full loss of Dll4, but not hemizygous loss of Dll4. (d) The 7 genes significantly deregulated after hemizygous loss of Dll4 are also deregulated after full loss of Dll4 signalling, but with very distinct amplitudes. (e) Myc target genes are upregulated with similar amplitude in coronary ECs with hemizygous and full loss of Dll4/Notch signalling. (f) Subepicardium venous ECs proliferate less (EdU+/ERG+) after the full loss of Dll4/Notch signalling (n=4 hearts per group). (g and h) Subepicardium venous ECs have increased ERK activation (P-ERK) and p21 expression after the full loss of Dll4/Notch signalling (n=3 hearts per group). (i) Analysis of EC proliferation in arterial and venous zones of E15.5 wildtype hearts (minimum n=4 hearts per group). (j and k) Analysis of EC proliferation in the arterial zone of control, Dll4^f/f-iDEC^ and Rbpj^f/f-iDEC^ mice (n=3 hearts per group). Scale bars, 100 um. Error bars indicate SD. *p < 0.05, ***p < 0.001. NS, non-significant.

Dll4-Notch signalling has also been implicated in the regulation of endothelial sprouting of retinal ECs ^8, 39, 45^. However, unlike retinal vessels, angiogenic coronary vessels with loss of Dll4/Notch signalling showed no clear increase in the expression of genes related with sprouting angiogenesis (Fig. 3i) or tip-cell enriched genes ^46, 47^ (Fig. 3j). This different molecular profile is consistent with the absence of filopodia or morphologically distinct sprouting cells in most coronary vessels of control or *Dll4* ^*f/f-iDEC-Pdgfb-24h*^ hearts at E14.5 (Fig. 3k). In coronary vessels, Dll4–Notch signalling thus appears to control genetic signatures related to endothelial proliferation and arteriovenous differentiation but not endothelial sprouting.

### Arterial development failure after hemizygous loss of Dll4/Notch signalling is associated to cell proliferation and not cell differentiation defects

In addition to the RNAseq analysis of coronary ECs with full loss of Dll4-Notch signalling, we also performed RNAseq on coronary ECs with a 50% (2-fold) decrease in Dll4 expression for 24h *(Dll4*^*f/Wt-iDEC-Pdgfb-24h*^ or *Dll4*^*HET-iDEC-Pdgfb-24h*^). In contrast with coronary ECs with full loss of *Dll4* expression, the 50% decrease in Dll4 expression resulted in only 7 DEGs at a Benjamin-Hochsberg adjusted p-value < 0.05 (Fig. 4a). Interestingly, in the hemizygous mutant hearts, pre-arterial ECs also failed to form arteries (Fig. 4b), even though they did not lose their pre-arterial identity. In *Dll4*^*f/Wt-iDEC-Pdgfb-24h*^ ECs, only *Dll4* itself and *Cx40/Gja5* were significantly downregulated (around 2 fold), and the expression of all other artery-specific genes did not change significantly (Fig. 4c). Major coronary arteries can thus fail to develop in Dll4–Notch hemizygous mutants even when acquisition of the pre-arterial molecular identity is not compromised. Interestingly, ECs with homozygous and hemizygous loss of Dll4 expression showed a significant increase in *Myc* (Figure 4d) and many of its downstream target genes^48, 49^ (Fig. 4e). Myc is one of the most important and ubiquitous regulators of cell proliferation and biosynthetic pathways ^50, 51^, including in ECs ^49^. Given the very small DEG set in *Dll4* ^*f/Wt-iDEC-Pdgfb-24h*^ coronary ECs, this data suggests that the Myc pathway is one of the most tightly repressed pathways by Notch signalling in coronary ECs.

### Notch inhibition results in opposing proliferative effects on venous and arterial coronary plexus

The data above show that ECs with full or hemizygous loss of Dll4-Notch signaling upregulate pathways linked to cell metabolism and proliferation, which should result in increased EC proliferation. Paradoxically, however, sub-epicardial venous ECs proliferate significantly less when they lose Dll4-Notch signalling or the transcription factor Rbpj (Fig. 4f and Extended Data Fig. 4c-j).

This result is consistent with the recently identified bell-shaped Notch dose-dependent regulation of EC proliferation, according to which ECs become arrested when exposed to highly mitogenic stimulation, induced by full loss of Notch signaling in a context of high VEGF signaling ^8^. Growing coronary vessels are exposed to high levels of VEGF secreted by the adjacent myocardium ^52^, resulting in ERK activation and coronary EC proliferation ^53^. In the full absence of Dll4-Notch signalling, P-ERK levels increased significantly in angiogenic coronary vessels, correlating with increased expression of the cell-cycle inhibitor p21 (Fig. 4g, h), which arrests angiogenesis ^8^.

Unlike the highly proliferative sub-epicardial venous vessels, arterial zone capillaries, located deeper in the myocardium, have higher Notch signalling and proliferate significantly less (Fig. 4i), as also reported previously ^27, 32^. EC proliferation in these arterial capillaries was increased upon homozygous or hemizygous loss of Dll4–Notch signaling or loss of Rbpj (Fig. 4j,k). This dual and location-dependent role of Notch is highly conserved, and also occurs in the retinal angiogenesis system, where angiogenic front ECs become arrested after inhibition of Dll4–Notch signaling, while periarterial and more mature ECs re-enter the cell cycle and proliferate more ^8^. Therefore, given that *Dll4*^*f/Wt-iDEC*^ ECs fail to develop arteries but do not lose their arterial identity, the only common factor in the arterial defects of *Dll4*^*f/f-iDEC*^ and *Dll4*^*f/Wt-iDEC*^ mutants is an increase in pre-arterial EC proliferation and the expression of genes related to the promotion of the cell cycle and metabolism.

### Repression of Myc by Notch and VEGF during arterialization

Our results suggest that the transcriptional Notch program induced in pre-arterial capillaries is associated with cell-cycle exit induced by lower P-ERK or Myc levels. However, VEGF, the most important endothelial promitogenic factor, is known to be required for arterialization in embryos and coronary artery development ^10, 52^. To investigate the influence of VEGFR2 and ERK signaling levels on arterialization, we analyzed coronary arterialization in *iMb-Vegfr2-Mosaic* mice ^33^, which allow the induction of a Cre-dependent genetic mosaic of cells with high (MbTomato+) or low (MbYFP+) levels of VEGFR2–ERK signaling (Fig. 5a-c). MbYFP+ (Vegfr2^low^) ECs were rarely found in the intramyocardial pre-arterial coronary plexus and did not form arteries, but they formed the endocardium and heart-surface venous vessels (Fig. 5b-g), likely due to genetic compensation by VEGFC-VEGFR3 signaling in this heart region ^54^. In contrast, MbTomato+ (Vegfr2^high^) ECs frequently formed arteries but tended not to incorporate in veins (Fig. 5e-g). These results show that ECs with very high VEGFR2–ERK signaling are more likely to form arterial vessels, contrasting our previous finding that Notch induces arterialization by suppressing ERK signaling and mitogenic activity. However, both Vegfr2/ERK overactivation and Notch activation induce cell-cycle exit during coronary angiogenesis (Fig. 5h,i), through a process dependent on a bell-shaped dose-response to mitogenic stimulation, according to which both low or very high ERK signalling levels induce cell-cycle exit, as previously identified in the retina angiogenesis system (Pontes-Quero et al., 2019). Low VEGFR2–ERK signaling also induced frequent cell-cycle exit (Fig. 5i), but this was insufficient to induce effective arterial differentiation, since MbYFP+ (Vegfr2^low^) cells could form endocardium but were not prone to form arteries (Fig. 5f,g). We reasoned that this likely reflects the requirement of VEGFR2 signaling for cell mobilization or adequate activation of Dll4 and Notch signaling in pre-arterial ECs ^10, 55–57^, and that Notch, not ERK signaling, subsequently mediates adequate suppression of the downstream Myc program. Supporting this interpretation, MbTomato+ (Vegfr2^high^) cells had high levels of ERK signaling (Fig. 5b,c), but relatively low levels of Myc (Fig. 5j,k), whereas ECs with loss of Dll4 or Rbpj signalling had high activity of both ERK (Fig. 4g) and Myc (Fig. 3e-h and Fig. 5l,m). These results indicate that is the low Myc levels induced by high arterial VEGF/Notch activity, that sets the cell-cycle or metabolic suppression that enables the subsequent arterial development.

**Fig. 5.**
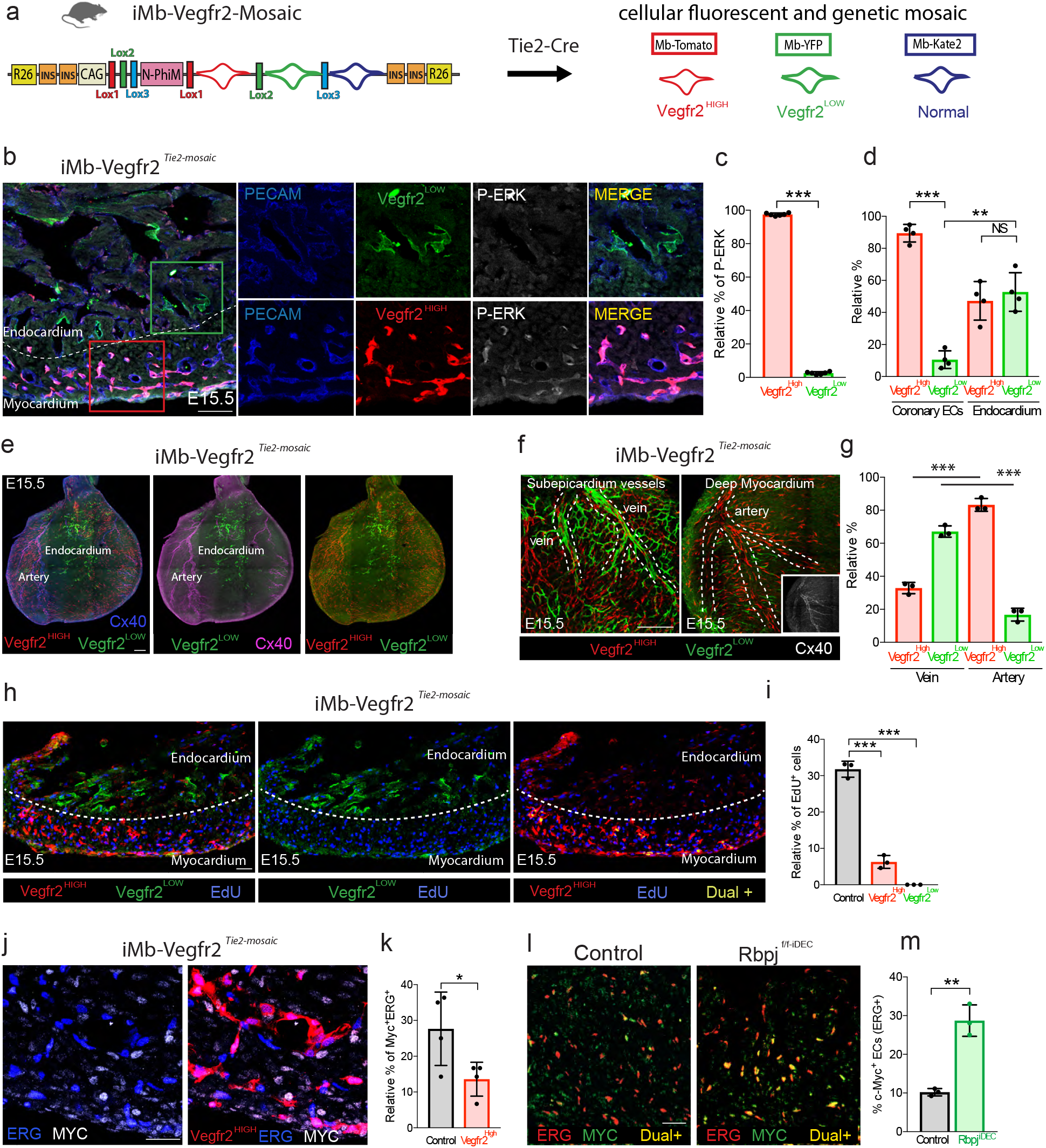
High VEGF/ERK signalling induces arterialization by suppressing EC proliferation. (a) Mice containing an iMb-Vegfr2-Mosaic allele were interbred with Tie2-Cre mice to generate a mosaic of coronary ECs having distinct VEGFR2 signalling levels. (b-e) Relative P-ERK levels and localization of Tomato+ (Vegfr2 high) and YFP+ (Vegfr2 low) in coronary vessels and endocardium (n=4 hearts per group). (f and g) Relative contribution of ECs with distinct Vegfr2/ERK signalling levels to coronary veins and arteries (n=3 hearts per group). Inset shows high Cx40 in Tomato+ arteries. (h and i) Analysis of proliferation (EdU+) in ECs with distinct Vegfr2/ERK signalling levels (n=3 hearts per group). (j and k) Comparison of Myc expression in Wildtype (Tomato−) and Vegfr2 high (Tomato+) ERG+ ECs (n=4 hearts per group). (l and m) Comparison of Myc expression in Wildtype and Rbpj mutant ERG+ ECs (n=3 hearts per group). Scale bars, 100 um. Error bars indicate SD. *p < 0.05, **p < 0.01, ***p < 0.001. NS, non-significant.

### Myc is a negative regulator of arterial differentiation

To define the role of Myc in coronary angiogenesis, we induced *Myc* deletion in ECs between E12.5 and E15.5. As expected, Myc was required for EC proliferation and coronary vessel growth, particularly in venous and periarterial capillaries, but less so in the arterial remodeling zone or the artery itself (Fig. 6a, b). The decrease in EC proliferation in *Myc*^*f/ft-iDEC*^ mutants ultimately delayed embryonic development. Some *Myc*^*f/ft-iDEC*^ mutant hearts were underdeveloped and lacked arteries (Extended Data Fig. 5a,b), whereas others had minor arterialization defects (Fig. 6a), presumably due to a lower Myc deletion efficiency. To elucidate the cell-autonomous role of Myc in arterial differentiation; without impacting the whole embryo and heart vascular development, we used the inducible genetic mosaic system (*iFlp*^*Tom-Cre/Yfp*^) presented above. By generating mice carrying the alleles *Apln-FlpO*, *iFlp*^*Tom-Cre/Yfp*^ *Mosaic* and *Myc*^*f/f*^ we were able to induce *Myc* loss-of-function genetic mosaics in Apln+ sinus-venosus-derived coronary ECs, which are progenitors of most coronary vessels and arteries ^58^. Although ECs with loss of Myc (Tomato+) proliferated less, they formed arteries at a much higher rate than the control (YFP+) cells (Fig. 6c). Thus, by enhancing the proliferative and metabolic activity of coronary ECs, Myc negatively regulates pre-arterial specification. These results are consistent with recent single-cell RNAseq data showing a significant decrease in *Myc* expression during coronary arterial differentiation ^23^. The significant decrease in *Myc* expression and function caused by the higher Notch activity in arterial vessels may explain why Myc deletion does not affect EC proliferation in the arterial zone (Fig. 6b).

**Fig. 6.**
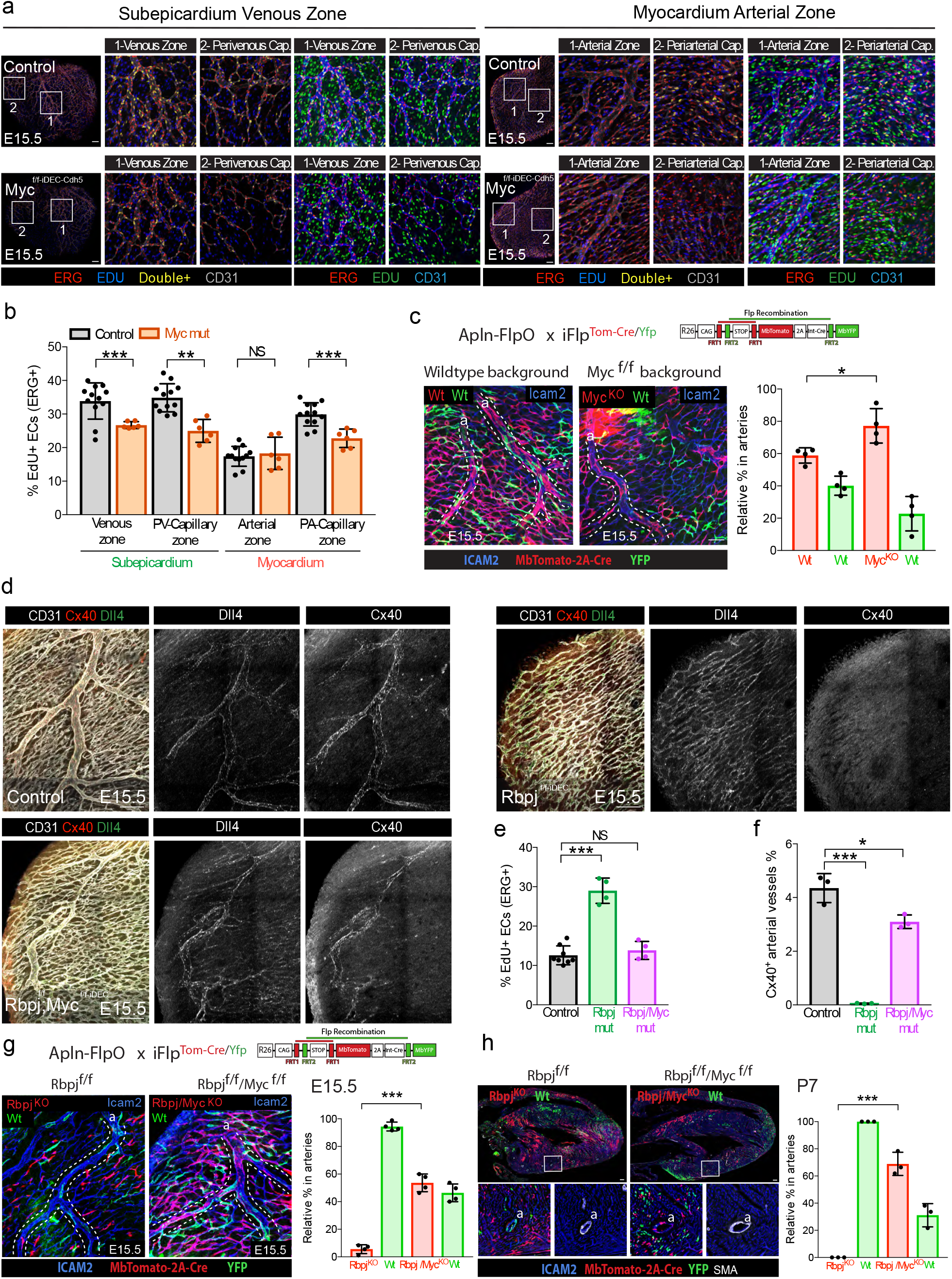
Arterial development can occur in the absence of Dll4 or Rbpj signalling when Myc function is suppressed. (a and b) EC proliferation is reduced in Myc^iDEC^ mutants, except in coronary vessels located in the arterial zone (minimum n=3 hearts per group). (c) Mice carrying the alleles Apln-FlpO and iFlp^MTomato-Cre/MYfp^ have induction of a mosaic of ECs in growing coronary vessels. When these alleles are on a Myc^fl/fl^ background Tomato-2A-Cre+ cells having Myc deletion (see also Extended Data Fig.1j,k and Extended Data Fig. 6e) are more likely to form arteries, despite their reduced proliferation rate (n=4 hearts per group). (d-f) Loss of Myc rescues the proliferation (ERG+/EdU+, see Extended Data Fig. 6d for chart e data) and arterial differentiation (Cx40) defects induced by the full loss of Rbpj (minimum n=3 hearts per group). (g and h) Mice carrying the alleles Apln-FlpO and iFlp^MbTomato-Cre/MYfp^ have induction of a mosaic of ECs in growing coronary vessels. When these alleles are on a Rbpj^fl/fl^ background Tomato-2A-Cre+ cells rarely form arteries, however in the Rbpj/Myc^fl/fl^ background they can form arteries at E15.5 (n=4 hearts per group) and P7 (n=3 hearts per group). Scale bars, 100 um. Error bars indicate SD. *p < 0.05, **p < 0.01, ***p < 0.001.

### Myc loss enables arterial development in the complete absence of the Notch-Rbpj transcriptional activator complex

Given the data presented above, we hypothesized that Notch inhibition interferes with arterial development by inducing Myc upregulation and cell-cycle progression in pre-arterial vessels, and not by directly inhibiting arterial differentiation. To test this hypothesis, we compared coronary EC proliferation and arterial differentiation in control embryos (littermates lacking the CreERT2 allele) and, *Dll4*^*f/Wt-iDEC*^ and *Myc*^*f/f*^*/Dll4*^*f/Wt-iDEC*^ embryos. *Myc* deletion was sufficient to revert the increase in proliferation and lack of arterialization caused by deleting a single copy of *Dll4* (Extended Data Fig. 6a-c).

Numerous studies have shown that the full loss of Dll4, Notch or Rbpj impairs arterial differentiation and arterialization in zebrafish and mice ^9, 10, 15–17^. So far, no other genetic mechanism has been shown to overrule or compensate for the absence of Notch signalling during arterialization. Given the data presented above, we analysed coronary arterialization in hearts combining full loss of *Rbpj* and *Myc*. Full deletion of *Myc* rescued the severe arterialization and arterial identity defects observed in hearts with full deletion of *Rbpj*. In contrast to *Rbpj*^*f/f-iDEC*^ mutants, *Rbpj*^*f/f*^ *Myc*^*f/f – iDEC*^ pre-arterial ECs are able to form large Cx40+ arteries at E15.5 (Fig. 6d-f and Extended Data Fig. 6d).

We next wanted to investigate whether *Rbpj*^*f/f*^ *Myc*^*f/f – iDEC*^ ECs could form functional arteries for longer periods; however, these mutants have global vascular defects incompatible with long-term embryo development. We therefore generated mice carrying the alleles *Apln-FlpO* (X-chromosome located and mosaic), *iFlp*^*MTomato-Cre/MYfp*^, *Rbpj*^*f/f*^ and *Myc*^*f/f*^, so that we could induce trackable dual *Myc/Rbpj* loss-of-function genetic mosaics in Apln+ sinus-venosus derived coronary ECs (Extended Data Fig. 6e). These vascular mosaic mutant mice developed normally and reach adulthood (Extended Data Fig. 6f). Unlike cells with single loss of *Rbpj*, most ECs with full loss of *Rbpj* and *Myc* were able to contribute to artery formation at E15.5 (Fig. 6g) and were still present in a high proportion in adult coronary arteries (Fig. 6h), indicating that they can maintain arterial identity and function for long periods.

These results suggest that Notch is a master regulator of arterialization not because it directly induces the arterial genetic program, but because it suppresses Myc and its downstream metabolic and cell-cycle activity, a prerequisite for subsequent acquisition of the coronary arterial cell fate.

## Discussion

Gene expression analysis studies have revealed that arterial and venous ECs express different sets of genes ^14, 23, 59–61^. Functional genetic studies showed that the loss of some of these artery or vein-specific genes leads to the specific impairment of arterial or venous development ^1, 9, 10, 15, 61–63^. However, a cell’s final identity or fate is often established after several transition steps, and establishing primary gene function in these consecutive steps is often difficult given the fast pace of biological processes and the simultaneous integration of many genetic and biophysical factors.

Activation of Notch signalling culminates in the transcriptional activation of multiple arterial-enriched genes, and is required for capillary ECs to form arteries ^9, 10, 15–17, 19^. This led to the conclusion that Notch induces the direct genetic differentiation or pre-determination of pre-arterial ECs. However, we are not aware of any studies showing the direct binding of the Notch downstream effector transcription factors Rbpj or Hey to the promoters or enhancers of arterial or venous genes. In addition, Notch overactivation *in vitro* fails to induce the expression of most arterial-specific genes ^59^.

More refined studies using the latest technologies in lineage tracing and imaging suggested that apart from the role of Notch in identity priming or pre-determination, ECs with high Notch are particularly able to migrate against the flow towards the arteries. This process allows the growth of distal arterial branches, a process involving a combination of Cxcl12-Cxcr4 chemokine signalling and an arterial fate-inductive stimulus ^21, 22, 64^. Here, we show that the sole inhibition of the Myc-dependent cell-cycle or biosynthetic activity by Notch, results in the adoption of an arterial phenotype, without the need for the Notch-dependent transcriptional cellular pre-determination or differentiation. Given the interplay and often mutually exclusive nature of cell proliferation and migration ^8^, it is reasonable to think that timely inhibition of the cell cycle by the higher Notch activity in pre-arterial ECs will render them more responsive to chemokine signalling or other cell migration cues. This in turn may facilitate their assembly and incorporation into developing arteries, adopting subsequently their inductive microenvironment and final arterial identity (Extended Data Fig.7).

This model is supported by several lines of evidence. Our genetic lineage tracing revealed biases in arteriovenous segregation of cells with distinct VEGF or Notch signaling levels; nevertheless, ECs with high Notch activity can form veins and ECs with low Notch activity (DN-Maml1+) or lacking the transcription factor Rbpj can form arteries. This would not be possible if a Notch-driven hard-wired transcriptional prespecification existed. In addition, the complete loss of Notch–Rbpj in already developed and mature arteries does not compromise their arterial phenotype or identity ^37, 65^. All the available evidence suggests that Notch function seems to be important only in capillary-to-arterial transition, a process associated with reduced metabolic and cell-cycle activity ^23,27, 32^. Our study shows that this transition can occur in the absence of Notch signaling so long as Myc function is shut off. The fact that capillary ECs can differentiate and form arterial vessels without the activity of the NICD-Rbpj transcriptional activator complex, indicates that this pathway does not directly prime ECs for arterial differentiation. Our results instead suggest that pre-arterial capillary ECs need Notch to inhibit Myc-dependent cell cycle and metabolism in order to prime them for subsequent arterial differentiation or mobilization. This is supported by the finding that Myc-depleted ECs tend to form more arteries than adjacent wildtype cells. Our results also explain how VEGF, a potent inducer of ERK and mitogenic activity, can induce arterial differentiation while also suppressing the cell cycle. A relatively mild increase in ERK activity, such as after the loss of Notch signaling, is known to induce Myc activity ^66, 67^; however, strong VEGF–ERK stimulation induces high Dll4–Notch signaling, and this reduces Myc levels and EC proliferation, inducing arterialization. The model proposed here for the regulation of arterialization also matches the recently published observation that overexpression of the venous-determining transcription factor CoupTFII–Nr2f2 in prearterial ECs compromises arterial development by inducing the expression of cell-cycle genes ^23, 68^.

From a translational standpoint, is known that atherosclerosis is an artery-specific disease, with no incidence in veins, and that coronary artery bypass surgery is more likely to fail when the graft is a heterologous saphenous vein rather than an artery ^69, 70^. Induction of collateral arterial growth is also seen as a promising new approach against cardiac ischemia caused by coronary artery disease ^71^. In this context, the ability to induce or reprogram the arterial or venous identities of established and quiescent vessels is of great interest. The mechanistic insights presented here advance knowledge of the process of arterialization and may enable better induction of this process during tissue growth, regeneration or ischemic cardiovascular disease.

## Methods

### Mice

The following mouse alleles were used: *iChr-Notch-Mosaic* ^33^; *iMb-Vegfr2-Mosaic* ^33^; *Tie2-Cre* ^34^, *PAC-Cdh5-CreERT2* ^72^; *Pdgfb-iCre-ERT2-IresEGFP* ^44^; *Nfact1-Cre* ^52^, *Dll4*^*flox /flox73*^, *Rbpj*^*flox/flox* 40^, *Myc*^*flox/flox* 74^. The *ROSA26*^*LSL-MbTomato-2A-H2B-GFP-2A-N1ICDP*^ mice were generated by targeting the construct indicated in Fig. 2a (full plasmid sequence and DNA to be deposited at Addgene # reference) to the Rosa26 locus of mouse ES cells with G4 background following standard procedures ^33^. The *iFlp*^*Tom-Cre/Yfp*^*Mosaic* mice were generated by targeting the construct indicated in Fig. 1i and Extended Data Fig. 1j (full plasmid sequence and DNA to be deposited at Addgene # reference) to the Rosa26 locus using the same procedure (To be described in detail elsewhere). The Apln-FlpO mice were generated by targeting via CRISPR/Cas9 assisted gene targeting (guide RNA target sequence GAATCTGAGGCTCTGCGTGC-AGG) in mouse G4 ES cells the HA-NLS-FlpO-WPRE-Sv40-pA-LoxP-PGK-Neo-polyA-LoxP cassette in frame with the ATG of the endogenous *Apln* gene (full plasmid sequence and DNA to be deposited at Addgene # reference). All primer sequences required to genotype this mice are provided in Extended Data Table 1.

To induce CreERT2 activity, 20mg tamoxifen (Sigma-Aldrich, T5648) and 10mg progesterone (Sigma-Aldrich, P0130) were first dissolved in 140ul absolute ethanol and then 860ul corn oil (20mg/ml tamoxifen and 10mg/ml progesterone). This solution was given via oral gavage (200ul, total dose of 4mg tamoxifen per mouse). Dosing and dissection schedules for individual experiments were as follows. For *N1ICDP* inducible gain-of-function experiments, tamoxifen was administered at E11.5 and E12.5, followed by dissection at E15.5 or E16.5 as indicated. For *Rbpj, Dll4* ^wt/fl^, *Myc* inducible loss-of-function experiments, tamoxifen was administered at E12.5, followed by dissection at E15.5. For *Dll4*^*wt/fl*^ and *Dll4*^*fl/fl*^ inducible short-term loss-of-function experiments, tamoxifen was administered at E13.5, followed by dissection at E14.5 (24h LOF, used for the RNAseq analysis). All mouse husbandry and experimentation was conducted using protocols approved by local animal ethics committees and authorities (Comunidad Autónoma de Madrid and Universidad Autónoma de Madrid—CAM-PROEX 177/14 and CAM-PROEX 167/17).

### Whole-mount immunofluorescence and 3D imaging of embryonic hearts

Whole embryos were dissected between E12.5 and E15.5, and their chests were fixed in 4% paraformaldehyde (PFA) on ice for 2h (E12.5) or 5h (E14.5-E15.5) with shaking. Chests were washed with shaking for 2×10min in PBS at room temperature (RT). Hearts were removed and cut on the ventral side (for artery morphological analysis) and the dorsal side (for venous morphological analysis), and the endocardium was trimmed with microdissection scissors (FST NO.15001-08, FST No.11210-10). For immunostaining, hearts were first blocked and permeabilized in PBBT solution (PBS with 0.5% Triton X-100 and 10% donkey serum) for 2h at RT with shaking. Subsequently, primary antibodies were added to fresh PBBT solution (see Key Resources Table in STAR Methods), and hearts were incubated at 4°C overnight with shaking. The next day, hearts were washed in PBT (PBS with 0.5% Triton X-100) for 6×15 min to remove unbound primary antibody. Secondary antibodies (see Key Resources Table in STAR Methods) were diluted in PBT, and samples were incubated at 4°C overnight with shaking, followed the next day by 6×15min washes in PBT. Heart samples were placed in the middle of a 100 micron spacer (Invitrogen, S24737) and mounted in Fluoromount-G (SouthernBiotech, Cat: 0100-01). Large Z-volumes of the samples were imaged using different objectives and tile scanning on a Leica SP5, Leica SP8 STED or Leica SP8 Navigator microscopes. Images were processed and analyzed with ImageJ/Fiji, Photoshop and Illustrator.

### Immunofluorescence on cryosections

Embryonic hearts were isolated and fixed in 4%PFA for 2h on ice and then washed with PBS 3×5min. After washing, hearts were incubated in 30% sucrose in PBS overnight at 4°C with shaking. Hearts were then embedded in OCT (Sakura), snap frozen on dry ice and stored at −80°C. Cryosections (25um) were cut with a Leica Cryostat (Leica CM1950). For immunofluorescence, slides were dried at RT for 10 min followed by washing with PBS 3×5min, then blocked with PBBT (0.5% Triton X-100 and 10% donkey serum in PBS) for 1h at RT and incubated with primary antibodies diluted in PBBT overnight at 4°C in a humidified chamber. The following day, slides were washed in PBS for 3×5min and then incubated with secondary antibodies diluted in PBS for 2h at RT in humidified chamber. Next, slides were washed in PBS at RT 3×5min and mounted in Fluoromount-G (SouthernBiotech, Cat: 0100-01) before confocal imaging.

For phosphoextracellular signal-regulated kinase (P-ERK) staining, we used a different protocol with the tyramide signal amplification kit (TSA Plus Fluorescein System; PerkinElmer Life, NEL774). Briefly, cryosections were dried at RT for 30min followed by washes in PBS 3×5min. Cryosections were then incubated in sub-boiling sodium citrate buffer (10mM, pH 6.0) for 30min, allowed to cool at RT for 30min, and incubated for 30 minutes in 3% H_2_O_2_ in methanol to quench endogenous peroxidase activity. Slides were rinsed in double-distilled (dd) H_2_O and washed in PBS 3×5min, followed by blocking in PBBT for 1h and incubation with the primary antibody in PBBT (see Key Resources Table in STAR Methods) overnight at 4°C. Slides were then washed and incubated for 2h with anti-rabbit-HRP secondary antibody at RT; after washes, the signal was amplified using the TSA fluorescein kit (NEL774).

### Immunofluorescence on paraffin sections

To detect the short-lived, and difficult to detect, cleaved/activated N1ICD epitope, we used the TSA amplification kit (NEL774) procedure in paraffin sections after antigen retrieval. Briefly, sections were dewaxed and rehydrated followed by antigen retrieval in sub-boiling sodium citrate buffer (10mM, PH=6.0) for 30min. The slides were cooled down at RT for 30min, followed by incubation for 30 minutes in 3% H_2_O_2_ in methanol to quench the endogenous peroxidase activity. Next, slides were rinsed in ddH_2_O and washed in PBS 3×5min, followed by blocking in PBBT for 1h, and incubation in PBBT with the primary antibody (see Extended Data Table 1) overnight at 4°C. Next, the slides were washed and incubated for 2h in anti-rabbit-HRP secondary antibody at RT, after washing, the signal was amplified using the TSA fluorescein kit (NEL774).

### Endothelial cell isolation, RNA extraction and qRT-PCR

Coronary ECs with distinct Notch signalling levels were isolated according to their fluorescence from *iChr2-Notch-Mosaic* or *ROSA26*^*LSL-MbTomato-2A-H2B-GFP-2A-N1ICDP*^ mice after crossing with *Tie2-Cre* mice. Fluorescent (mutant) and non-fluorescent (wildtype/control) cells were isolated as indicated further below. To isolate and profile coronary ECs with unaltered or half or full loss of Dll4 signalling, we intercrossed *Pdgfb-iCre-ERT2-EGFP* mice with wildtype females or *Dll4*^*flox/flox*^ *Pdgfb-iCre-ERT2-EGFP* male mice with wildtype or *Dll4* ^*flox/flox*^ females. Tamoxifen was administered at E13.5 and embryos collected at E14.5. For this experiment, three groups of hearts (each group contained 6 to 9 embryos) with EGFP expression were obtained: controls, Dll4 hemizygous mutants, and Dll4 homozygous mutants.

Embryonic hearts were dissected, minced and digested for 20min at 37°C with 2.5 mg/ml collagenase type I (Thermofisher), 2.5 mg/ml dispase II (Thermofisher) and 50 μg/ml DNAseI (Roche). Dissociated samples were filtered through a 70-μm cell strainer, and cells were centrifuged (300g, 4°C for 5 min). Cell pellets were gently resuspended in blood lysis buffer (0.15 M NH_4_Cl, 0.01M KHCO_3_ and 0.01 M EDTA in distilled water) and incubated for 10 minutes on ice to remove erythroid cells. Cells were centrifuged (300g at, 4°C for 5 min), and cell pellets were gently resuspended in blocking solution (DPBS without Ca^2+^ or Mg^2+^ and containing 3% dialyzed FBS (Termofisher)) and incubated at 4°C with shaking for 20min. Cells were centrifuged (300g at 4°C for 5 min), resuspended and incubated for 30min at 4C with APC rat-anti-mouse CD31 (BD Pharmigen, 551262, 1:200). Cells were then centrifuged (300g, 4°C for 5 min), resuspended and washed in DPBS without Ca^2+^ or Mg^2+^ and centrifuged again, and cell pellets were resuspended in blocking solution. Cells were kept on ice until used for FACS. DAPI (5 mg/ml) was added to the cells immediately before FACS. Cells were sorted in a FACS Aria Cell Sorter (BD Biosciences). For each group, approximately 2000-4000 DAPI negative APC-CD31+ cells without or with fluorescence (GFP or Tomato or Cherry or Cerulean) were sorted directly to buffer RLT (RNAeasy Micro kit - Qiagen) and RNA extracted according to the manufacturer instructions and stored at −80°C. This RNA was later used for qRT-PCR or RNAseq analysis.

For quantitative real time PCR (qRT-PCR), total RNA was retrotranscribed with the High Capacity cDNA Reverse Transcription Kit with RNase Inhibitor (Thermo fisher, 4368814). cDNA was preamplified with Taqman PreAmp Master Mix containing a Taqman Assay-based pre-amplification pool containing a mix of the following Taqman assays (*Cdh5* (Mm00486938_m1), *Hey1*(Mm00468865_m1), *Hey2*(Mm00469280_m1), *Hes1*(Mm01342805_m1), *Dll4*(Mm00444619_m1), *Rbpj*(Mm01217627_g1), *Myc*(Mm00487804_m1). Preamplified cDNA was used for qRT-PCR with the same gene specific Taqman Assays and Taqman Universal Master Mix in a AB7900 thermocycler (Applied Biosystems).

### Next Generation Sequencing

NGS services were provided by the Genomics Unit of CNIC. 1ul of stored RNA per sample was quantified using the Agilent 2100 Bioanalyzer RNA6000 pico total RNA chip. After quantification, 700pg of total RNA were used for cDNA amplification with the SMART-Seq v4 Ultra Low Input RNA Kit (Clontech-Takara). 1 ng of amplified cDNA was used to generate barcoded libraries using the Nextera XT DNA library preparation kit (Illumina). In brief, cDNA was fragmented and adapters are added in a single reaction followed by amplification and clean-up steps. Library size was checked using the Agilent 2100 Bioanalyzer High Sensitivity DNA chip, and library concentration was determined with a Qubit® fluorometer (ThermoFisher Scientific). Libraries were sequenced on a HiSeq2500 (Illumina) to generate 100 bases paired-end reads.

### RNA-seq data analysis

RNA-seq data were analyzed by the CNIC Bioinformatics Unit. Sequencing reads were processed with a pipeline that used FastQC (Babraham Bioinformatics - https://www.bioinformatics.babraham.ac.uk/projects/fastqc/raham.ac.uk/projects/fastqc/) to evaluate their quality, and cutadapt^75^ to trim sequencing reads, thus eliminating Illumina and SMARTer adaptor remains, and discard reads shorter than 30 bp. Resulting reads were mapped against mouse transcriptome GRCm38.76, and gene expression levels were estimated with RSEM^76^. Expression count matrices were then processed with an analysis pipeline that used the Bioconductor package limma^77^ for normalization (using the TMM method) and differential expression testing; matrix processing considered only genes expressed with at least 1 count per million (CPM) in at least as many samples as the condition with the least number of replicates. Three pairwise contrasts were performed: *Dll4*^*f/f-iDEC-Pdgfb-24h*^ (abbreviated in raw data file as KO) vs *Control*^*iDEC-Pdgfb-24h*^ (abbreviated in raw data file as Wt) and *Dll4*^*f/wt-iDEC-Pdgfb-24h*^ (abbreviated in raw data file as Het) vs *Control*^*iDEC-Pdgfb-24h*^. Changes in gene expression were considered significant if associated to Benjamini and Hochberg adjusted p-value < 0.05. Complementary enrichment analyses with GSEA ^78^ were performed for each contrast, using the whole collection of genes detected as expressed (12,872 genes) to identify gene sets that had a tendency to be more expressed in either of the conditions being compared. Gene sets, representing pathways or functional categories from the Hallmark, KEGG, Reactome and Biocarta databases, and GO collections from the Biological Process, Molecular Function and Cellular Component ontologies were retrieved from MsigDB ^79^. Enriched gene sets with FDR < 25% were considered of interest.

### Ultrasound analysis of cardiac function

Mice aged 2 to 5 months were analysed for cardiac function at the CNIC Advanced Imaging Unit. Briefly, mice were anesthetized by inhalation of isoflurane/oxygen and examined with a 30-MHz transthoracic echocardiography probe and a Vevo 2100 ultrasound system (VisualSonics, Toronto, Canada). End-systolic and end-diastolic LV volumes (LVESV and LVEDD), ejection fraction (EF) and fractional shortening (FS) were calculated from short-axis and long-axis B-mode views.

### In vivo EdU labelling and EC proliferation detection

To detect EC proliferation rates in embryonic hearts, 50 μg/g body weight EdU (Invitrogen— A10044) was injected intraperitoneally into pregnant mice 1 h before dissection. Embryonic hearts were isolated for cryosection or wholemount analysis. EdU signals were detected with the Click-it EdU Alexa Fluor 647 or 488 Imaging Kit (Invitrogen, C10340 or C10337). Briefly, after all other primary and secondary antibody incubations, samples were washed according to the immunofluorescence staining procedure and then incubated with Click-iT EdU reaction cocktails for 40 min, followed by DAPI counterstaining.

### Statistical analysis

Two groups of samples with a Gaussian distribution were compared by unpaired two-tailed Student *t*-test. Comparisons among more than two groups were made by ANOVA followed by the Turkey pairwise comparison. Graphs depict mean +/-SD as indicated, and differences were considered significant at p < 0.05. All calculations were done in Excel and final datapoints were analysed and represented with GraphPad Prism. No randomization or blinding was used, and animals or tissues were selected for analysis based on their genotype, the detected Cre-dependent recombination frequency, and the quality of multiplex immunostaining. Sample size was chosen according to the observed statistical variation and published protocols.

## Data availability

The RNAseq raw data will be deposited in a public repository. The source data underlying all figures and charts will be provided as a supplementary source data file. All other data supporting the study findings are available from the corresponding author upon request. This includes additional raw data such as unprocessed original pictures and independent replicates, which are not displayed in the manuscript but are included in the data analysis in the form of graphs.

## Author Contributions

W.L. and R.B. designed experiments and interpreted results. W.L. executed the vast majority of the experiments. I.G.G engineered the DNA constructs and validated the *iFlpMosaic* and *Apln-FlpO* mice. M.F.C. supported the RNAseq, FACS and qRT-PCR analysis. V.C.G. gave general technical assistance and established some of the ES cell lines used to generate mouse models. R.B. and W.L. wrote the manuscript.

## Competing Interests

The authors declare no competing interests.

## Acknowledgements

Research was supported by the European Research Council (ERC-2014-StG - 638028), the CNIC Intramural Grant Program – Severo Ochoa (11-2016-IGP-SEV-2015-0505), and the Ministerio de Economia, Industria y Competitividade y Ministerio de Ciencia y Innovación (MEIC: SAF2013-44329-P, RYC-2013-13209 and MCIN: SAF2017-89299-P). The CNIC is currently supported by the Ministerio de Ciencia e Innovación (MCIN) and the Pro CNIC Foundation and is a Severo Ochoa Center of Excellence (SEV-2015-0505). Wen Luo received a Marie Curie FP7 COFUND CNIC International Postdoctoral fellowship. Macarena Fernández-Chacón and Irene García González were supported by PhD fellowships from Fundación La Caixa (CX_E-2015-01 and CX-SO-16-1, respectively).

We thank Simon Bartlett and Susana Rocha for English editing; José Luis de La Pompa and Donal Macgrogan for scientific input throughout the project; all members of the CNIC Transgenesis units for the support in the generation of the mouse lines; all members of the CNIC Microscopy Unit for their input in confocal imaging and the CNIC Genomics and Bioinformatic Units for the RNAseq analysis. We also thank J.L. Pompa, F. Radtke, R.H. Adams, M. Torres and T. Honjo for sharing the *Tie2-Cre, Dll4* ^*floxed*^,*Cdh5(PAC)-CreERT2, Mycf*^*loxed*^ and *Rbpj*^*floxed*^ mice, respectively.

---

**Extended Data Fig.1.**
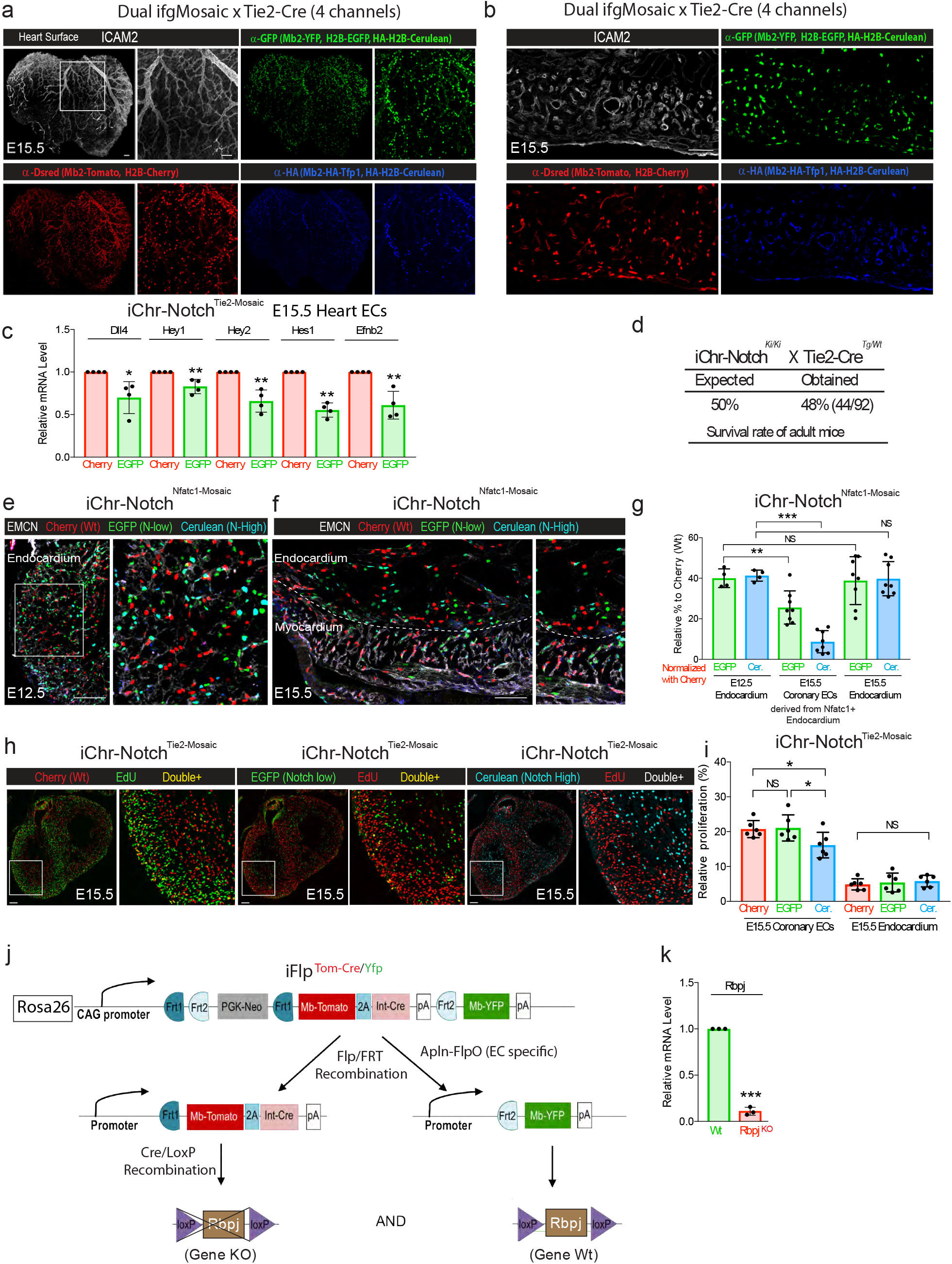
Multispectral genetic mosaics to analyse the proliferation and fate of heart ECs. (a and b) Individual channels of multispectral confocal imaging of the heart surface coronary vasculature and section showed in main Fig. 1a. (c) qRT-PCR analysis of Notch target genes in H2B-GFP+/DN-Maml1+ and H2B-Cherry+ (wildtype) Heart ECs collected by FACS (n=4 pooled litters). (d) Survival rate of adult mice containing the iChr-Notch-Mosaic and Tie2-cre alleles follows the expected mendelian frequencies indicating no lethality. (e and f) Confocal images of N1ICDP^Nfatc1-Cre^ embryonic hearts sections showing the localization and relative frequency of ECs having distinct Notch signalling levels. (g) Quantification of experiments represented in e and f showing how the relative percentage of each type of cell changes throughout development according to their location (n=4 hearts per stage). (h and i) Analysis of cell proliferation in Cherry+ and GFP+/DN-Maml1+ ECs (n=3 hearts per group). (j) Illustration showing the iFlp^Tom-Cre/Yfp^ Mosaic allele and expected outcomes in cells expressing Flp recombinase. (k) qRT-PCR analysis of Notch pathway genes in CD31+/MbYFP+ and CD31+/MbTomato+ ECs collected by FACS from Apln-FlpO iFlp^Tom-Cre/Yfp^ Rbpj^f/f^ E15.5 hearts (n=3 pooled litters). Residual Rbpj expression may reflect contamination of the MbTomato+ sample with RNA from wildtype cells as shown in Fernandez-Chacon et al., 2019. Scale bars, 100 um. Error bars indicate SD. *p < 0.05, **p < 0.01, ***p < 0.001. NS, non-significant.

**Extended Data Fig.2.**
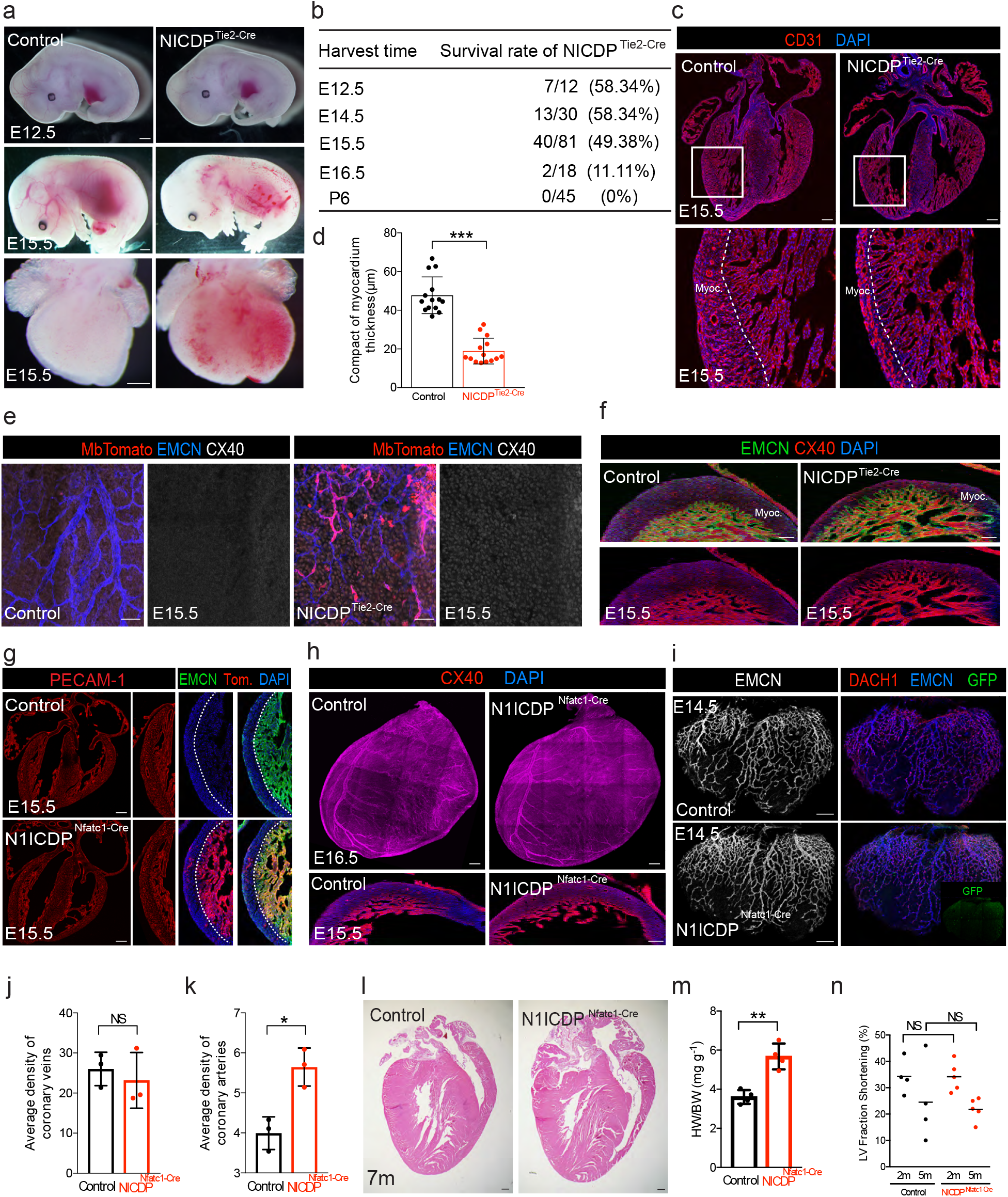
Overactivation of Notch in coronary ECs, not in endocardium, compromises coronary vessel and heart development. (a) Brightfield images of E12.5 and E15.5 embryos and hearts showing appearance of microcirculatory defects at E15.5. (b) Statistical analysis of mice survival rate per stage indicates that most mutant mice do not pass embryonic stage E16.5. (c-d) CD31 staining of control and N1ICDP^Tie2-Cre^ heart sections showing vascular defects and thinner myocardial wall (inset) in mutant hearts (n=7 hearts per group). (e) Confocal images showing immunostaining of the heart surface venous plexus for the arterial marker Cx40. ECs with Notch activation (Tomato+) do not express Cx40 and form veins. (f) Confocal images of heart sections showing an increase in Cx40 immunosignals in the myocardium (see also chart in Fig. 2p). (g) Induction of N1ICDP expression (Tomato+) in the Nfatc1+ endocardium and its lineages does not cause major cardiovascular defects at E15.5. (h-k) Comparative analysis of coronary arteries (h) and veins (i) development in control and N1ICDP^Nfatc1-Cre^ hearts (n=3 hearts per group). (l) Hematoxylin and Eosin staining of 7 months-old hearts of control and N1ICDPNfatc1-Cre mice. Mutant hearts are larger. (m and n) Quantification of heart weight and LV fraction shortnening (n=4 animals per group). Scale bars, 100 um. Error bars indicate SD. *p < 0.05, **p < 0.01, ***p < 0.001. NS, not significant.

**Extended Data Fig. 3.**
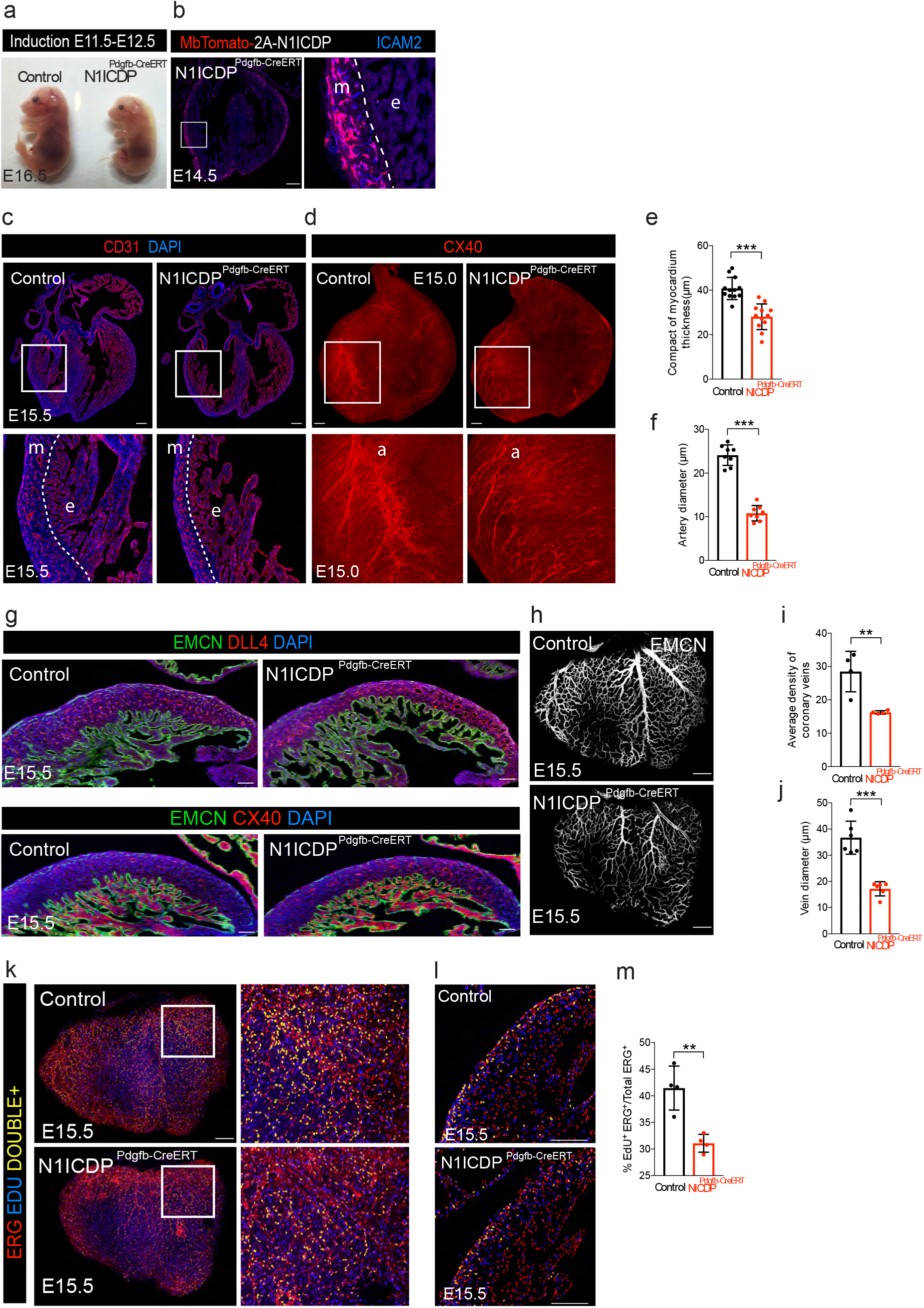
Induction of Notch activity in N1ICDP^Pdgfb-CreERT^ myocardial vessels results in similar cardiovascular defects to N1ICDP-^Tie2-Cre^ hearts. (a) Stereomicroscope analysis of mutant embryos and control littermates showing growth retardation at E16.5. (b) Immunostaining for the reporter MbTomato shows the induction of the allele in Pdgfb+ myocardial (m) vessels (Icam2+) and not endocardium (e). (c and e) Thinner myocardial wall and vascular defects in N1ICDP^Pdgfb-CreERT^ hearts (n=4 hearts per group). (d and f) Reduced coronary arterial diameter in N1ICDP^Pdgfb-CreERT^ hearts (n=4 hearts per group). (g) Expression of the arterial proteins Dll4 and Cx40 in control and N1ICDPPdgfb-CreERT hearts. (h-j) Reduced coronary vessel density and vein diameter in N1ICDP^Pdgfb-CreERT^ hearts (n=3 hearts per group). (k-m) Reduced proliferation (ERG+/EdU+) of N1ICDP^Pdgfb-CreERT^ subepicardium venous vessels (k) and myocardial vessels (l, m). (n=4 hearts per group). Scale bars, 100 um. Error bars indicate SD. **p < 0.01, ***p < 0.001.

**Extended Data Fig. 4.**
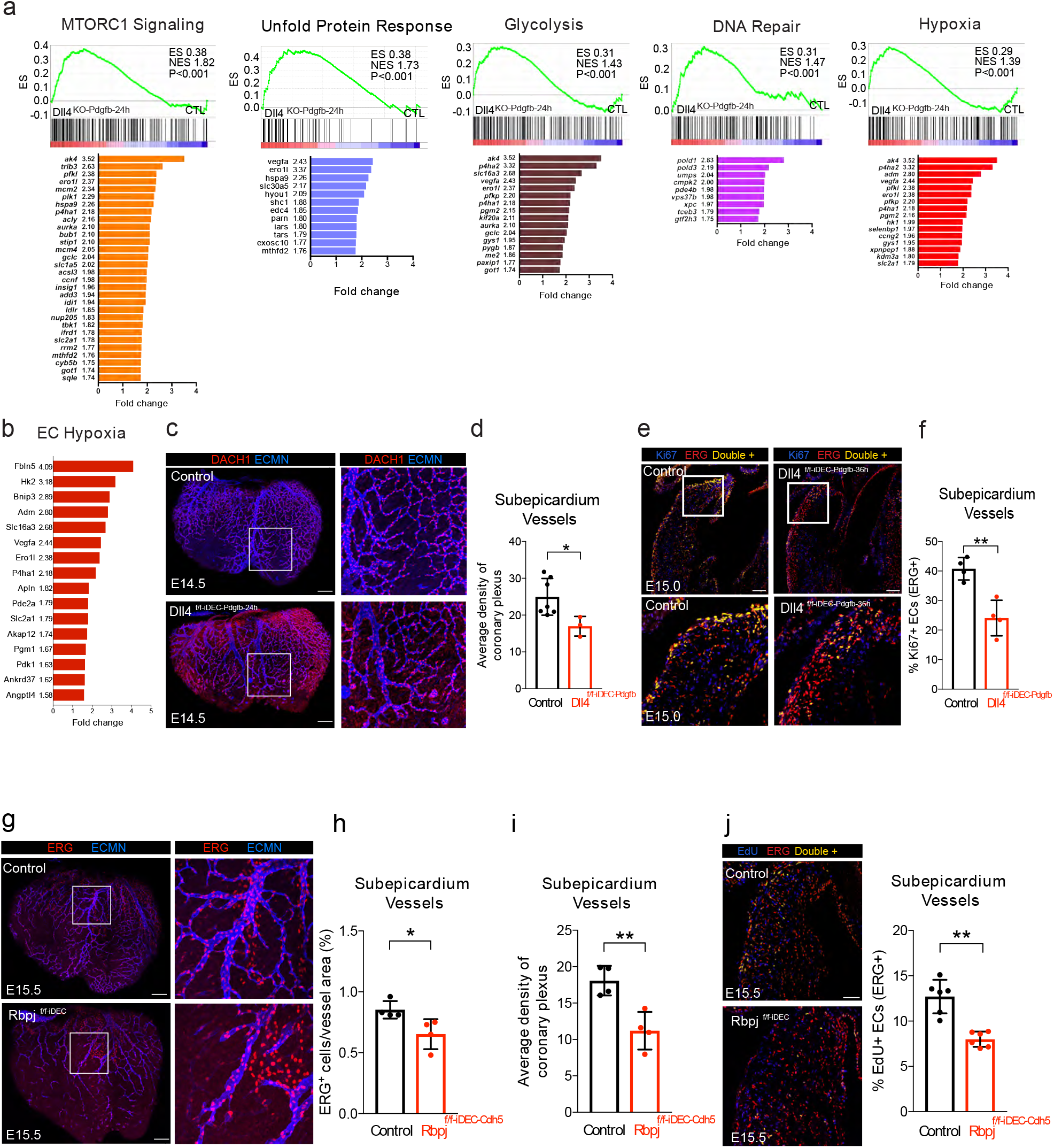
Induction of proliferation and metabolic pathways after Dll4 deletion does not result in more subepicardial EC proliferation. (a) Complete list of deregulated genes within a selected enriched gene set (GSEA). (b) Selected list of genes shown before to be upregulated when HUVECs are exposed to hypoxia 72, 73. (c and d) Dll4 deletion in Pdgfb+ myocardial ECs for 24h reduces the average density of heart subepicardial vessels (n=3 hearts per group). (e and f) Dll4 deletion for 36h reduces the frequency of Ki67+ ECs in subepicardium vessels (n=4 hearts per group). (g-i) Rbpj deletion reduces the number of ECs and average density of subepicardium vessels (n=4 hearts per group). (j) Rbpj deletion reduces the frequency of proliferation (EdU+/ERG+) of subepicardium vessels (n=3 hearts per group). Scale bars, 100 um. Error bars indicate SD. *p < 0.05, **p < 0.01.

**Extended Data Fig. 5.**
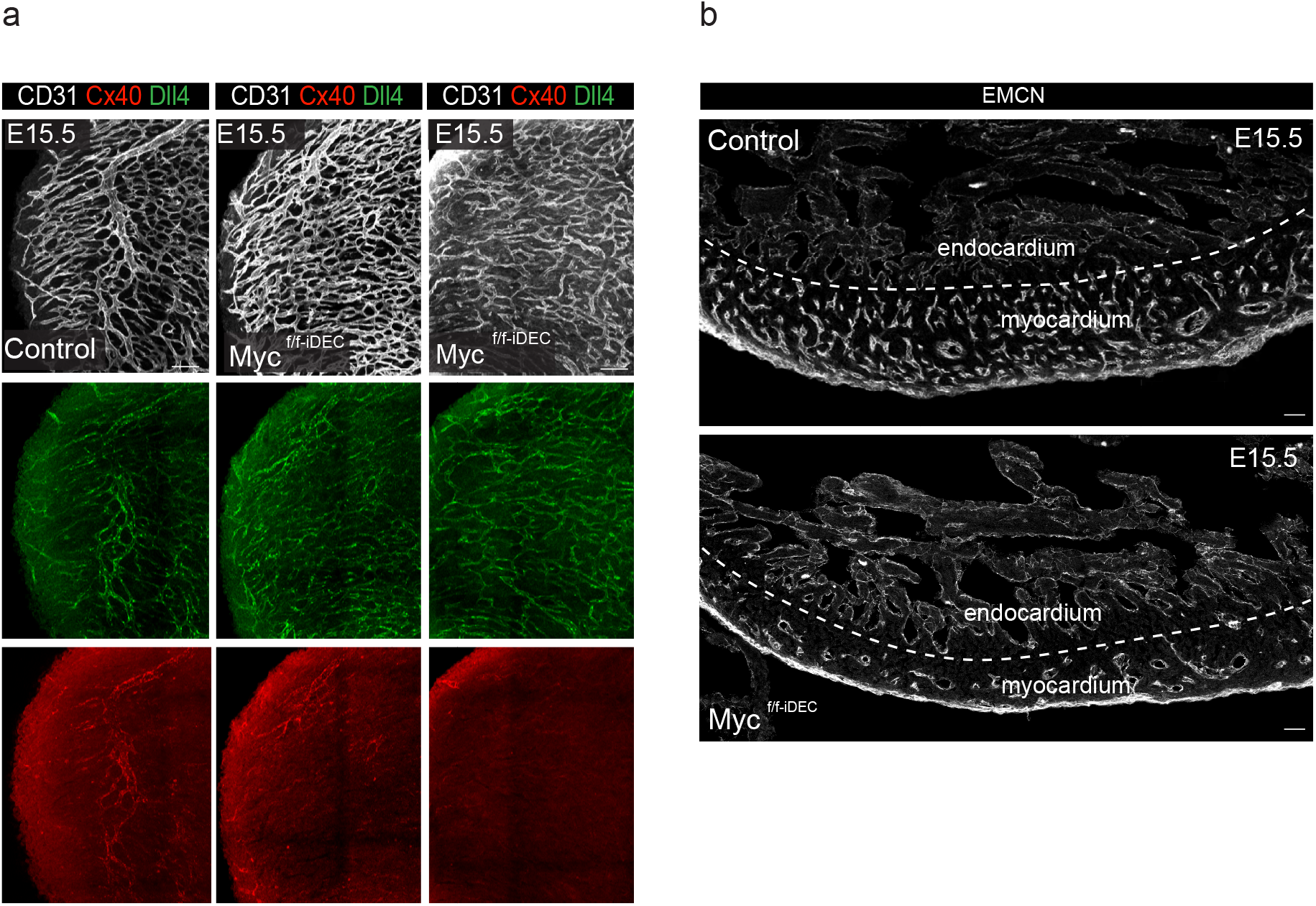
Global Myc deletion compromises cardiovascular development. (a) Wholemount analysis of control and different Myc mutant hearts showing that global Myc deletion compromises the development of coronary vessels and consequently arteries (Cx40+). Note that efficiency of Myc deletion may be variable among mutant littermates. (b) Sectional analysis of control and Myc mutant hearts showing a reduction in the number of coronary vessels in the myocardium which causes a reduction in its thickness. Scale bars, 100 um.

**Extended Data Fig. 6.**
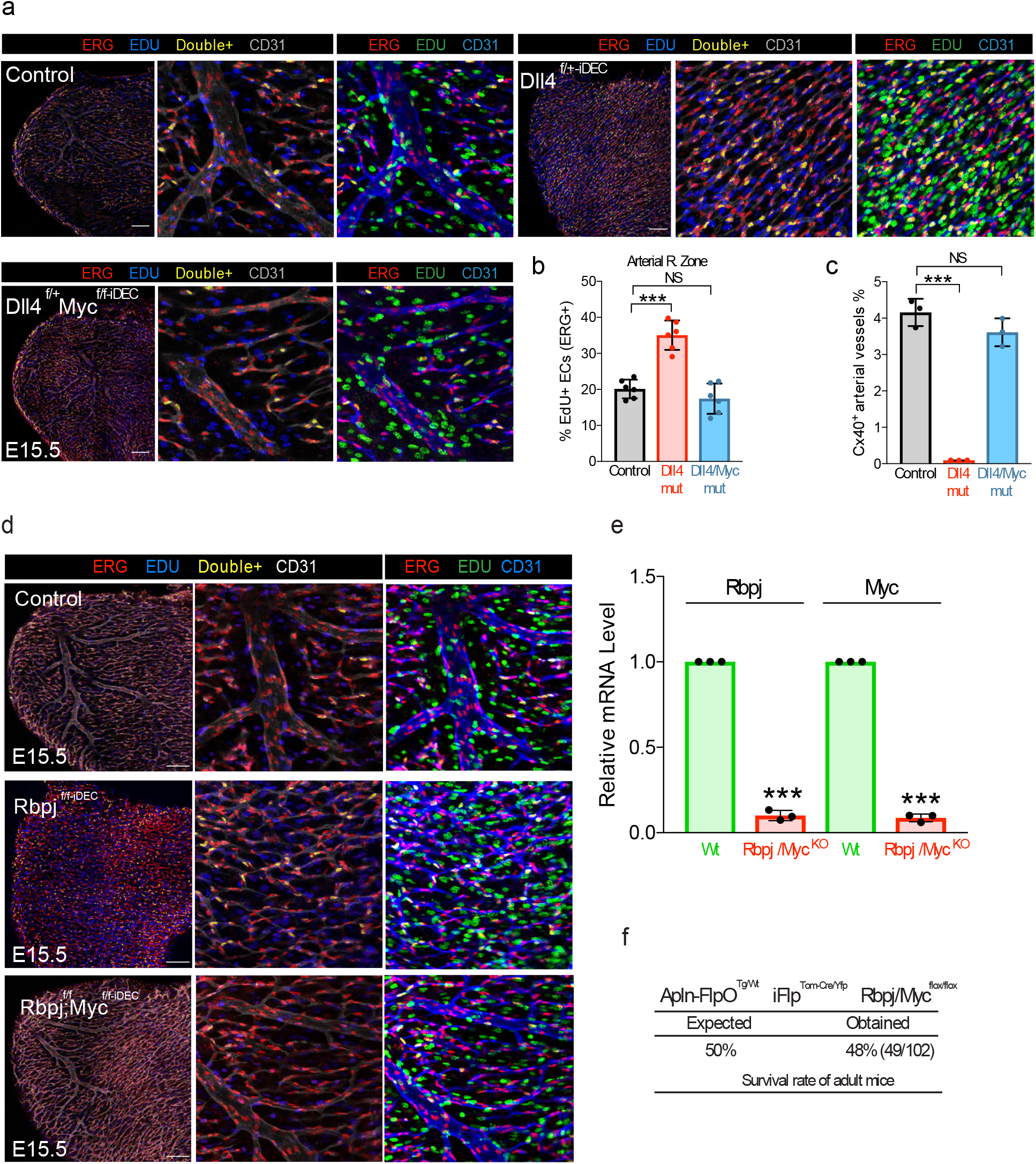
Loss of Myc rescues the proliferation and arterialization defects caused by the loss of Dll4 or Rbpj. (a-c) Loss of Myc rescues the proliferation (ERG+/EdU+) and arterial differentiation (Cx40) defects induced by hemizygous loss of Dll4 (minimum n=3 per group). (d) Confocal images of whole hearts showing proliferative ECs (ERG+/EdU+) in the arterial zone of control and mutant hearts (see Fig. 6e chart for quantification). (e) qRT-PCR analysis of Tomato+ and YFP+ cells isolated from Apln-FlpO iFlp^Tom-Cre/Yfp^ Rbpj^f/f^ Myc^f/f^ hearts showing the expected difference in Rbpj and Myc expression between the wildtype (YFP+) and mutant (Tomato+) cells. Residual Rbpj and Myc expression may reflect contamination of the MbTomato+ sample with RNA from wildtype cells as shown in Fernandez-Chacon et al., 2019. (f) Table showing that Apln-FlpO iFlp^Tom-Cre/Yfp^ Rbpj/Myc^f/f^mutant mice are found at the expected mendelian inheritance frequency. Scale bars, 100 um. Error bars indicate SD. ***p < 0.001.

**Extended Data Fig. 7.**
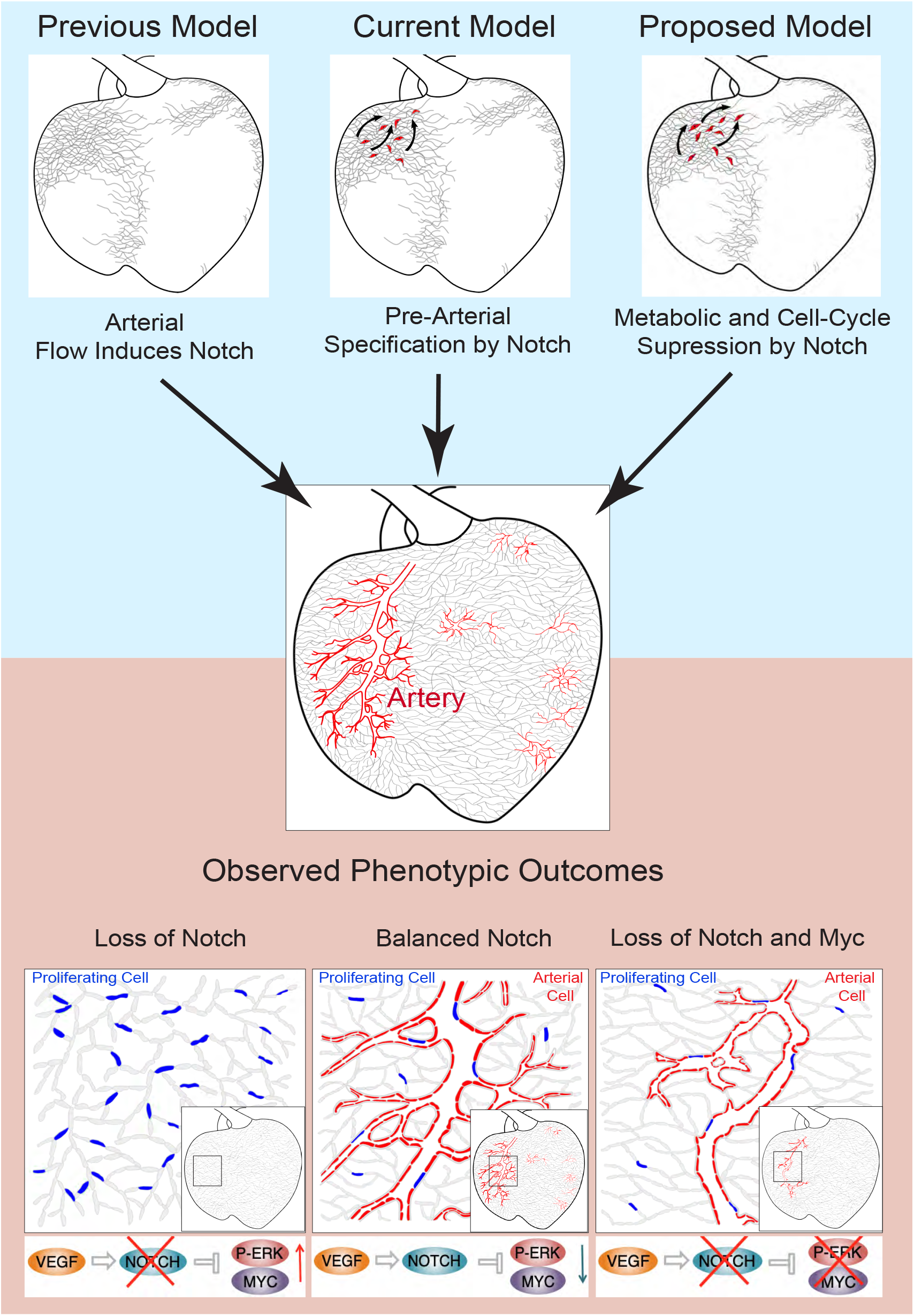
Models for the regulation of arterialization. The initial model used to describe arterialization was based on the assumption that an increase in pulsatile and oxygenated blood flow induces the direct differentiation and development of arterial vessels. The current model is based on the evidence that pre-arterial capillary ECs have higher Notch-Rbpj activity and higher expression of arterial genes. This model assumes that arterial fate is determined before the increase in arterial blood flow, through the direct induction of arterial gene expression via the Notch-Rbpj transcriptional complex. Here, we propose a model in which Notch-Rbpj suppresses Myc-dependent cell cycle or biosynthetic activity, rendering ECs more permissive to the adoption of an arterial phenotype without the need for Notch-dependent genetic pre-determination or differentiation. This model is supported by the observed phenotypic and transcriptomic outcomes in several mouse mutant models. In the absence of Notch, ERK and Myc activities are higher, and induce higher metabolic activity and proliferation, preventing subsequent arterial differentiation or mobilization. The arterial development defects caused by the loss of the Notch-Rbpj transcriptional activity in pre-arterial ECs can be reversed by decreasing Myc-dependent mitogenic and metabolic activities. These findings indicate that Notch controls arterialization by regulating the cell cycle and not differentiation.

**Extended Data Table 1.**
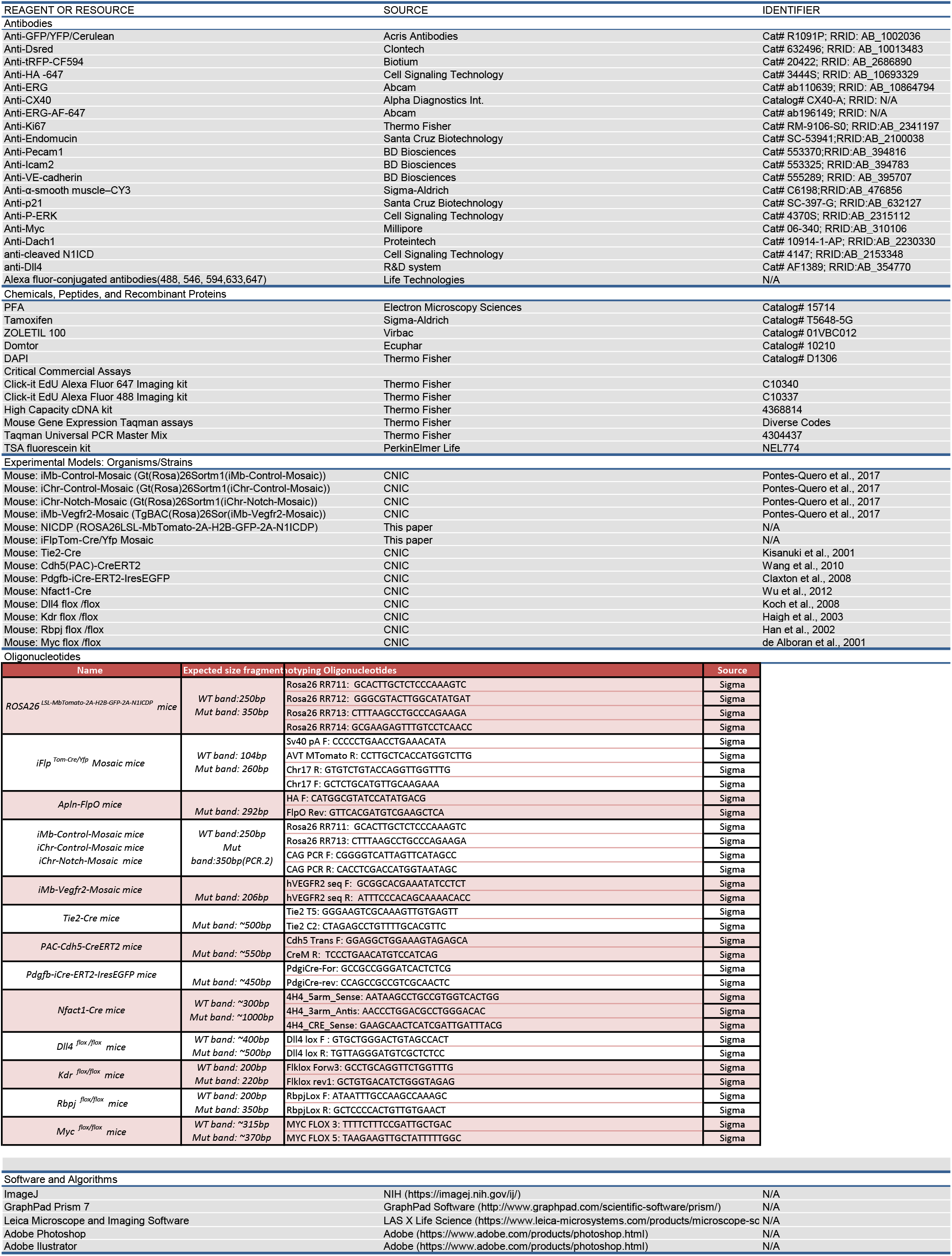

## Notes

### Competing Interest Statement

The authors have declared no competing interest.

